# The Role of the Sex Chromosomes in the Inheritance of Species Specific Traits of the Shape of the Copulatory Organ in Drosophila

**DOI:** 10.1101/161224

**Authors:** Alex M. Kulikov, Svetlana Yu. Sorokina, Anton I. Melnikov, Nick G. Gornostaev, Dmitriy G. Seleznev, Oleg E. Lazebny

**Affiliations:** Department of Evolutionary Genetics of Development, Koltzov Institute of Developmental Biology Russian Academy of Sciences, Moscow, 119334, Russia; Department of Ecology of Aquatic Invertebrates, Papanin Institute for Biology of Inland Waters Russian Academy of Sciences, Borok village, Yaroslavl Region, 152742, Russia

## Abstract

The sex chromosomes of the parental species, D. virilis and D. lummei were tested for the effect on trait dominance in the shape of the copulatory system in the interspecific crosses. The origin of the sex chromosome and the paternal genotype were found to affect the trait dominance in D. lummei × D. virilis progeny and backcross males heterozygous for the autosomes. A correlated variability analysis showed that the two sex chromosomes exert unidirectional effects, shifting dominance towards the conspecific phenotype. The effect of the X chromosome is to a great extent determined by epigenetic factors associated with the paternal genotype.

## INRODUCTION

Reproductive isolation contributes to the evolutionary process by allowing diverging species to accumulate genetic variation at adaptively important traits independently. Isolation is ensured by pre- and postzygotic isolating mechanisms, which differ in evolutionary origin and physiological basis. Postzygotic isolation arises as independent genetic variations accumulate in isolated populations and cause sterility and/or inviability of hybrid progenies because of non-additive interactions of the alleles at various loci, while each allele does not exert a deleterious effect in the gene pool of its original population (Bateson, 1909; Dobzhansky 1936, 1937a, b; Muller 1940, 1942). Prezygotic isolation results from the variation driven by sexual selection and prevents mating of individuals from different populations possessing different adaptations. In allopatric speciation, selection of traits acting at the copulation stage starts when postzygotic incompatibility has already formed. Reinforcement (Kirkpatrick, 2001; Schuler et al., 2016) and the sexual selection or sexual conflict (Trivers, 1972; Maynard Smith, 1976; Arnqvist, Nilsson, 2000; Johnstone, Keller, 2000) mechanisms can be involved in this case. In sympatric speciation, a prezygotic barrier should start forming before postzygotic isolating mechanisms are involved. It is noteworthy that experimental findings agree with the assumption that prezygotic isolation accumulates at a higher rate as compared with postzygotic isolation. Coyne and Orr (Coyne, Orr, 1989, 1997) have estimated the parameters in experiments with q of many closely related *Drosophila* species and confirmed a higher accumulation rate for prezygotic isolation.

To discuss the formation rate and genetic basis of the two types of isolating barriers, it is necessary to understand the similarities and differences of their underlying processes. Postzygotic isolation is neutral or random for proper isolation of related species. An independent accumulation of variation in different evolutionary lineages leads to fixation of the alleles that show incompatibility and, in the case of sex chromosome linkage, gives origin to Haldane’s rule (Turelli, Orr, 1995, 2000), which postulates a reduction in fitness components for the heterogametic sex (Haldane, 1922). Low dominance values at incompatibility loci are required for Haldane’s rule to occur concerning viability, while more complex interactions of dominance and sex-specific incompatibility are required for Haldane’s rule to occur for fertility (Turelli, Orr, 2000). The role that the sex chromosomes play as a main genetic structure element responsible for hybrid incompatibility is additionally illustrated by two other empiric regularities, which are known as the substantial X effect and sex-limited asymmetry of hybrid viability and fertility in reciprocal crosses (Coyne, Orr, 1989; Turelli, Moyle, 2007).

Prezygotic isolation results from a selection at sex-limited traits, mostly those associated with mating behavior. Compared with autosomes, the X and W chromosomes are enriched in sexually antagonistic genetic variation because alleles exerting alternative effects in different sexes are more likely to be fixed (Fry, 2010). X- and W-linked variation is consequently more likely to be targeted by selection at sex-limited traits.

To explain the role that the sex chromosomes play in the evolution of isolating mechanisms, it is essential to consider the effects of all three evolutionary factors: mutagenesis, selection, and genetic drift. Chromosome variation due to mutation depends on the total evolutionary time that a given chromosome spend in the male genome and shows different extreme magnitudes in the case of the sex chromosomes, being the highest in the Y and Z chromosomes and the lowest in the X and Z chromosomes (Kirkpatrick, Hall, 2004). Thus, the action of this evolutionary factor does not explain the substantial X effect but implies that the Y and Z chromosomes accumulate substantial genetic burden and degrade in the absence of recombination (Charlesworth, 1991). The conclusion is based on the assumption that all chromosomes accumulate variation due to mutation at the same rate in males. Studies of meiotic sex chromosome inactivation (MSCI) in males still make it possible to assume that heterochromatin forming in a major part of the sex chromosomes in pachytene (Turner, 2007) prevents repair mechanisms from fully restoring heterochromatin sequences, thus substantially increasing the mutation rate of the two sex chromosomes relative to that of the autosomes. If so, the circumstance provides a plausible explanation for the substantial X effect, faster-X theory (Coyne, Orr, 2004), and faster male theory (Wu, Davis, 1993).

In contrast, the significance of genetic drift is well explained by the chromosome population effective size, which is smaller in the sex chromosomes compared with autosomes (Mank et al., 2009). Alleles associated with pre- and postzygotic isolation are accordingly more likely to be fixed on the sex chromosomes under the influence of random genetic drift processes.

Selection differently affects sex chromosome variation during the formation of prezygotic vs. postzygotic isolation. The fixation of new alleles, which is essential for incompatibility to arise, implies the effect of positive selection only. Postzygotic incompatibilities are not selected directly, but their choice is mediated by functional associations of particular genes with viability and fertility traits. The sex chromosomes are enriched in sexually antagonistic genes (Gibson et al., 2002; Johnson, Lachance, 2012) and may more often be targeted by selection at differentially expressed traits. Genes involved in gametogenesis act as targets in the case of postzygotic isolation; and genes involved in mating behavior, in the case of prezygotic isolation. In species with XY sex determination, the X chromosome undergoes demasculinization (Sturgill et al., 2007; Johnson, Lachance, 2012), while postzygotic isolation mechanisms act more often in males, in agreement with Haldane’s rule. The dominance theory (Muller, 1942; Orr, 1993; Turelli, Orr, 1995) postulates that recessive variations of X-chromosomal incompatibility loci are expressed in hemizygous males and explains well a viability decrease unrestricted to sex-linked traits. The discrepancy between X-chromosome demasculinization and fertility impairment in males are possible to explain by nonadditive interactions of alleles and, primarily, recessive negative epistasis. Epistatic interactions normally ensure homogametic sex-limited expression and are altered in a hybrid genome, leading to aberrant expression in the heterogametic sex to cause infertility and a drop in fitness (Johnson, Lachance, 2012).

Following the drive hypothesis, selection may additionally act indirectly, by suppressing sex chromosome meiotic drive, which results from sex chromosome evolution and permanent internal conflict between sex determination systems (Tao, Hartl, 2003; Tao et al., 2007a,b; Meiklejohn, Tao, 2010). For example, various systems of sex chromosome segregation distorters have been identified in the X chromosome and autosomes in *Drosophila* species by an incompatibility locus analysis; the systems include pseudogenes, repeat regions (Tao et al., 2007a,b; Bayes, Malik, 2009), and heterochromatin blocks responsible for failure of sister X chromosomes to segregate during mitosis in the hybrid genome (Ferree, Barbash, 2009; Cattani, Presgraves, 2012).

The role that the Y chromosome plays in the selection-driven formation of prezygotic isolation depends to a substantial extent on the sex wherein a particular trait is expressed. Given that the X chromosome experiences demasculinization and a direct effect of positive selection, X-chromosome variation is mostly restricted to female choice traits (Bailey et al., 2011). However, demasculinization of the X chromosome is not absolute, and the hemizygous status of the X chromosome dramatically increases the effect of selection in males (Haldane, 1924; Kirkpatrick, Hall, 2004), while not excluding an accumulation of alleles associated with particular characteristics of male mating behavior. Asymmetry of male success in reciprocal interspecific crosses of related species (Throckmorton, 1982; Markow, 1981; Ehrman, Wasserman, 1987) formally resembles Haldane’s rule but differs in arising directly at the stage of choosing a partner from a related species, being based on the differences accumulated in the respective genomes. Unequal partner preferences in reciprocal crosses depend on the reaction norm of each of the species concerning partner choice traits (Arnold et al., 1996) and do not suggest linkage with the sex chromosomes.

In *Drosophila*, the X chromosome has been shown to play a role in several species-specific features that are observed in male mating behavior and prevent heterospecific mating, the set including various elements of the courting ritual and various means of signal generation and perception. When cuticular hydrocarbons (CHCs) are utilized as cues in *D. virilis* and *D. americana texana*, a leading role in choosing the mating partner is played by the male, which discontinues tapping and loses interest in the female if the female is heterospecific. The capabilities of recognizing the female type and discontinuing the courting ritual are associated mostly with the X chromosome (Nickel, Civetta, 2009). Sharma et al. (Sharma et al., 2012) have observed significant sex dimorphism in CHC profile distribution in isofemale *D. simulans* lines, thus confirming that the sex chromosomes are involved in regulating expression of the genes belonging to the metabolic cascades of hydrocarbon synthesis. Characteristics of the male mating song have been studied in crosses between closely related *Drosophila* species of the *sophophora*, *obscura*, and *virilis* groups (Hoikkala, Lumme, 1987; Suvanto et al., 1994; Williams et al., 2001; Huttunen, Aspi, 2003; Päällysaho et al., 2003; Gleason, Ritchie, 2004; Hoikkala et al., 2000, 2005). The results have shown that all chromosomes, including the sex chromosomes, contribute to the species-specific variation in mating song elements, while the set of chromosomes or loci associated with the differences vary among related species pairs. A role of the X chromosome has been demonstrated for the majority of the variants tested. Arbuthnott (Arbuthnott, 2009) has analyzed a large set of reproductively isolating traits with known genetic architectures and concluded that the majority (25 of 36) of the traits are controlled by several loci of relatively large effects. The conclusion was that the majority (70%) of traits responsible for intraspecific and interspecific differences are controlled by a few loci of relatively large effects. The variation due to interspecific differences displayed no predominant association with the sex chromosomes and substantially differed from intraspecific variation. Epistatic interactions were found to play a significant role in the intraspecific and interspecific variation of the male courtship song.

Copulation as a final stage of mating behavior determines the efficiency of insemination of the female and the contribution of the male genotype to the gene pool of the next generation. The copulation efficiency depends on the copulation duration and the insemination reaction. A crucial role in the latter is played by male accessory gland secreted proteins (Acps), which evolve at a far greater rate as compared with hemolymph proteins (Chen, Stumm-Zollinger; 1985; Chen, 1996; Wolfner, 1997; Nurminsky, 2005; Haerty et al., 2007; McGraw, Clark, 2008). A total of 46 genes are known to code for Acps in *D. melanogaster*, and only six of them are on the X chromosome. A decrease in female fertility in heterospecific crosses of *D. virilis* and *D. americana* is associated with three *D. americana* and two *D. virilis* recessive autosomal genes (Sweigart, 2010). The finding agrees with the idea that isolating barriers rapidly arise from a divergence of a few genes and demonstrates that the X chromosome is weakly involved in the formation of prezygotic isolating barriers.

The copulation duration directly depends on how well the male copulatory system matches the female genitalia in shape. Jagadeeshan and Singh (Jagadeeshan, Singh. 2006) have observed that in crosses between the closely related species *D. melanogaster* and *D. simulans*, mating behavior was directly associated with the species-specific shapes of the posterior lobes of the genital arch and the cercus, which are external structures of the *Drosophila* male copulatory system. The time course of copulation stages depended on the mechanic coupling of the male and female genitalia, and the coupling depended on the male external genital structures. The final stage of tight genital coupling was 2–5 times shortened in heterospecific crosses, while the preceding unstable coupling stage was prolonged. The findings indicate that female insemination may be incomplete in heterospecific crosses and that a particular mechanism exists to allow mating flies to be sensitive to the specificity of their contact. Laurie and colleagues have analyzed the genetic basis of species divergence to the shapes of male copulatory system elements in *Drosophila*. A series of studies was carried out with *D. simulans* and *D. mauritiana* to genetically analyze the species-specific distinctions in the shape of the epandrium, which is an external male genital structure, and at least eight linkage groups involved were localized to the X chromosome and chromosomes 2 and 3 (Liu J. et al., 1995; 1996; Laurie et al., 1997; Zeng et al., 2000). Additive interactions were mostly observed for the respective loci.

The shape of the male genitalia is the most rapidly evolving among all morphological characters, and its control system is a target of sexual selection (Eberhard, 1985). However, the shape of the epandrium, which is the posterior lobe of the genital arch, has actually been used as the only model to study the genetic basis of species-specific differences in the shape of the copulatory system in *D. simulans* and *D. mauritiana*. Do the X chromosome and autosomes play the same role in species-specific traits related to the shape of the copulatory system in other species groups? Does the Y chromosome contribute to the inheritance of these traits? In this study, we used interspecific crosses between *D. virilis* and *D. lummei* and backcrosses of F_1_ hybrid males with *D. virilis* females to evaluate the contribution of the sex chromosomes and the total contribution of the autosomes to the degree of dominance of the *D. virilis* and *D. lummei* phenotypes in traits related to the shape of the male copulatory system.

## MATERIALS AND METHODS

Strains *D. lummei* 200 (Serebryanyi Bor, Moscow) and *D. virilis* 160 were from the *Drosophila* collection of the Institute of Developmental Biology and carried the following recessive autosomal markers: *broken* (*b*) on chromosome 2, *gapped* (*gp*) on chromosome 3, *cardinal* (*cd*) on chromosome 4, *peach* (*pe*) on chromosome 5, and *glossy* (*gl*) on chromosome 6. The first two markers are expressed as gaps in the second transverse and L2 veins, respectively; the other markers determine different eye colors, which are well identifiable visually. F_1_ hybrid males were obtained from direct crosses of *D. virilis* females with *D. lummei* males and reciprocal crosses of *D. lummei* females with *D. virilis* males. In addition, F_b_ males were obtained by backcrossing hybrid males with *D. virilis* females. Crossing schemes are shown below.

1. Direct crosses P♀X_Vi_ X_Vi_, A_Vi_ A_Vi_ × ♂X_Lu_Y_Lu_, A_Lu_ A_Lu_ → F_1_ ♂X_Vi_ Y_Lu_, A_Vi_ A_Lu_ (abbreviated F_1_X_Vi_Y_Lu_) ♀X_Vi_ X_Vi_, A_Vi_ A_Vi_ × ♂F_1_ X_Vi_ Y_Lu_ A_Vi_ A_Lu_ → F_b_ ♂X_Vi_ Y _Lu_, A_Vi_ A_Lu_ (abbreviated F_b_X_Vi_Y_Lu_, A_H_); ♂X_Vi_ Y _Lu_, A_Vi_ A_Vi_ (abbreviated F_b_X_Vi_Y_Lu_, A_Vi_);
2. Reciprocal crosses P ♀ X_Lu_ X_Lu_, A_Lu_ A_Lu_ × ♂ X_Vi_ Y_Vi_, A_Vi_ A_Vi_ → F_1_ ♂X_Lu_ Y_Vi_, A_Vi_ A_Lu_ (abbreviated F_1_X_Lu_ Y_Vi_) ♀X_Vi_ X_Vi_, A_Vi_ A_Vi_ × ♂ F_1_ X_Lu_ Y_Vi_, A_Vi_ A_Lu_ → F_b_ ♂X_Vi_ Y_Vi_, A_Vi_ A_Lu_ (abbreviated F_b_X_Vi_Y_Vi_, A_H_); ♂X_Vi_ Y _Vi_, A_Vi_ A_Vi_ (abbreviated F_b_X_Vi_Y_Vi_, A_Vi_);

A_Vi_, A_Lu_, A_H_ are the *D. virilis*, *D. lummei*, and heterozygous autosomes, respectively.

Backcross males heterozygous at all autosomes and fully homozygous backcross males were selected for further analysis. Backcrosses with heterozygous F_1_ males were performed to exclude autosomal recombination events and to allow a rigorous analysis of the effects for variants of homozygous and heterozygous combinations of autosomes of the parental species.

All crosses were carried out at 25°C; a standard food medium and glass tubes of 22 mm in diameter were used; the progeny density was 50–70 flies per tube.

### Analysis of morphological structures

The mating organ was dissected using thin steel needles in a drop of water under a binocular microscope at a magnification of 12×8. To remove adipose structures preparations were incubated in boiling 10% NaOH.

A total of 136 preparations were examined, including 14 from *D. lummei* males, 13 from *D. virilis* males, 25 from F_1_ males obtained by crossing ♂ *D. lummei* × ♀ *D. virilis*, 31 from F_1_ males obtained by crossing ♂ *D. virilis* × ♀ *D. lummei*, 10 from F_b_ males with the genotype X_Vi_ Y_Vi_, A_Vi_ A_Lu_, 28 from F_b_ males with the genotype X_Vi_Y_Lu_, A_Vi_ A_Lu_, 5 from F_b_ males with the genotype X_Vi_Y_Vi_, A_Vi_ A_Vi_, and 10 from F_b_ males with the genotype X_Vi_Y_Lu_, A_Vi_ A_Vi_.

Morphometry was carried out using organ images, which were obtained using a Jen-100C electron microscope in the scanning mode at an accelerating voltage of 40 kV and an instrumental magnification of 300–500×. The sagittal view of the phallus was conventionally divided into four areas: the aedeagus body, gonites, apodeme, and cook. A coordinate grid was superimposed on the image. A conventional point at the junction of the aedeagus area, gonites, and apodema (referred to as the central point) was used as a landmark. Morphometric parameters (MPs) were obtained as distances between the intercrosses of coordinate lines with each other and the image outline. A scheme of measurements is shown in Fig. 1; MP characteristics are summarized in Table 1. The declinations of the cook and apodeme (axes MP 2 and MP 28) from axis MP 1 were expressed in radians and designated α and β, respectively.

**Table 1.** Morphometric parameters and their characteristics

| **MP** | **Morphometric characteristics** |
| --- | --- |
| **1** | Aedeagus length to the cook base |
| **2** | Maximum aedeagus length |
| **3** | Cook length |
| **4** | Distance from the MP1 axis to the lower outline as measured 93.75% of the MP1 length away from point 1 |
| **5** | Aedeagus height measured 93.75% of the MP1 length away from point 1 |
| **6** | Distance from the MP1 axis to the lower outline as measured 75% of the MP1 length away from point 1 |
| **7** | Aedeagus height measured 75% of the MP1 length away from point 1 |
| **8** | Distance from the MP1 axis to the lower outline as measured 50% of the MP1 length away from point 1 |
| **9** | Aedeagus height measured 50% of the MP1 length away from point 1 |
| **10** | Aedeagus height measured 25% of the MP1 length away from point 1 |
| **11** | Distance from the MP1 axis to the lower outline as measured 25% of the MP1 length away from point 1 |
| **12** | Aedeagus height at the base |
| **13** | Maximum distance from the MP1 axis to the upper outline |
| **14** | Distance from the aedeagus base to the projection of the highest point of the outline onto the MP1 axis |
| **15** | Distance from the MP1 axis to the elbow of the upper outline |
| **16** | Distance from point 1 to the projection of the elbow point of the upper outline onto the MP1 axis |
| **17** | Distance from the MP1 axis to the lowest point of the aedeagus outline |
| **18** | Distance from point 1 to the projection of the lowest point of the aedeagus outline onto the MP1 axis |
| **19** | Distance from the projection of the upmost point of the ventral part of the aedeagus outline to point 1 |
| **20** | Distance from the MP1 axis to the upmost point of the ventral part of the aedeagus outline |
| **21** | Gonite width at the base |
| **22** | Curve depth of the ventral part of the paramere |
| **23** | Distance from the MP1 axis to the paramere outline as measured at the base |
| **24** | Paramere length at the ventral curvature |
| **25** | Paramere length in the central part (at the MP23 midpoint) |
| **26** | Distance from the MP23 midpoint to the outline of the basal part of the paramere (characterizes the paramere bending angle) |
| **27** | Paramere height in the central part (at the MP26 midpoint) |
| **28** | Apodeme length |
| **29** | Apodeme width measured 25% of the MP28 length away from point 1 |
| **30** | Apodeme bending parameter (25% of the MP28 length away from point 1) |
| **31** | Apodeme width measured 50% of the MP28 length away from point 1 |
| **32** | Apodeme bending parameter (50% of the MP28 length away from point 1) |
| **33** | Apodeme width measured 75% of the MP28 length away from point 1 |
| **34** | Apodeme bending parameter (75% of the MP28 length away from point 1) |

**Fig. 1.**
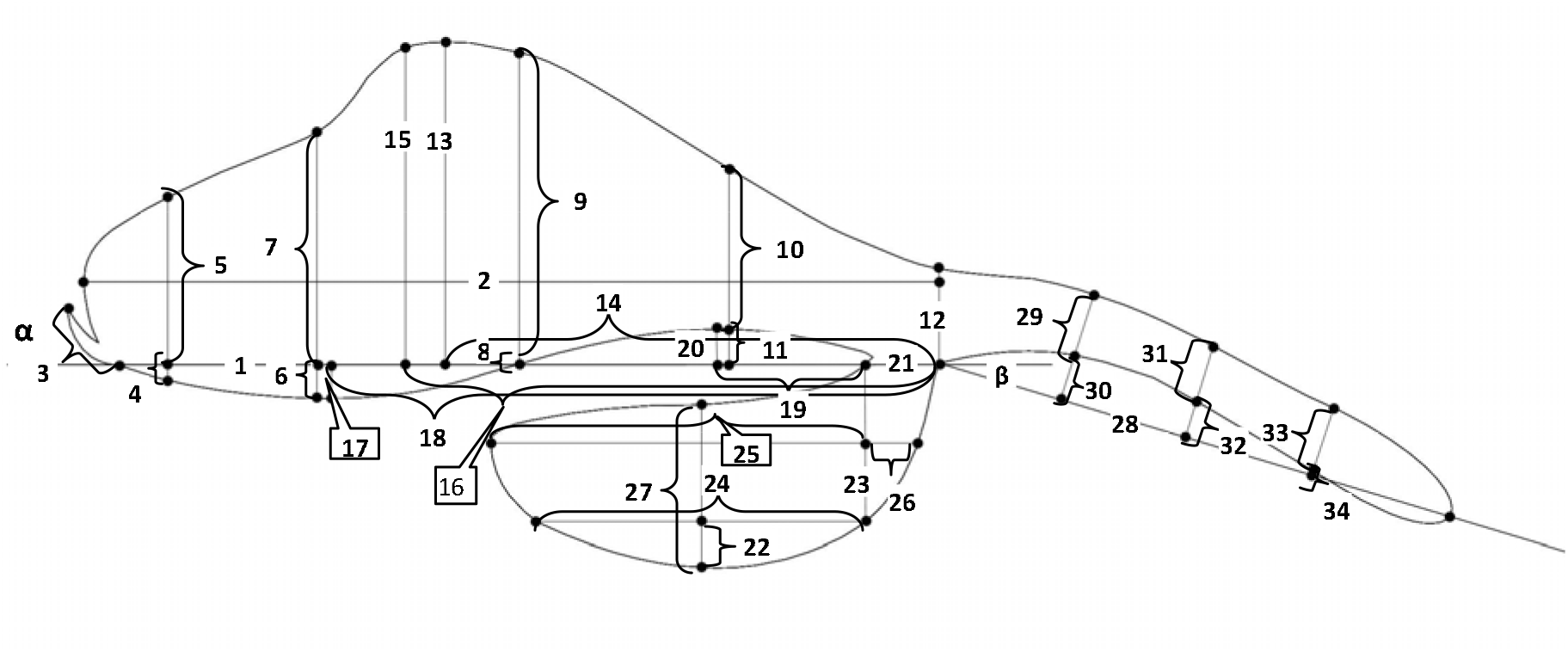
Scheme of morphometric parameters of the copulatory system measured in males of the *Drosophila* species of the *virilis* group.

To exclude the dimensionality factor, MP indices (MPIs) were calculated according to a method used previously to evaluate the inter- and intraspecific variations of the genitals in the virilis species group (1). MPIs were obtained as MP-to-MP 1 ratios and were numbered according to the numbers of respective MPs. The traits expressed in radians were included in the analyses without normalization.

### Statistical analyses

were carried out using the program IBM SPSS Statistics v. 23 and the R 3.3 statistical analysis environment with the packages “vegan”, “lmPerm”, “psych”, and “rcompanion”. Trait dominance in interspecific crosses was evaluated by comparing the trait variance in fly samples from the parental and hybrid strains; backcross samples included only males with a fully heterozygous autosome set. Differences were tested for significance by multivariate ANOVA; groups displaying homogeneity of variance were identified for each trait by post-hoc comparisons, using the Gabriel test and Tukey’s HSD test with the Bonferroni correction.

The effects of the sex chromosomes, autosomes, paternal genotype and their interactions on the traits were evaluated using samples from the parental, F_1_, and F_b_ populations, including both homozygotes and heterozygotes for the autosome set. A linear combination of weighted genetic factors was used as a model. The formal model is as follows:

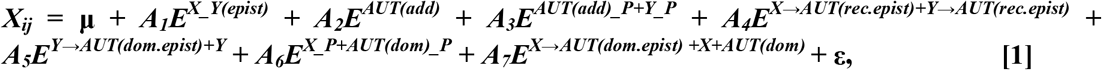

where *X*_*ij*_ is the trait value in the sample of a particular genotype; μ is the mean; ε is the fraction of variance unexplained; *A*_*n*_ values are the weights or indicator variables corresponding to the regression coefficients *E*^*n*^; and *n* indicates the respective genetic factor or factor combination: X and Y, independent effects of the respective sex chromosomes; AUT(add), the additive effect of recessive autosomal alleles; and AUG(dom), the dominant effect of the autosomes. Pairwise interactions of the factors were also considered, including X_Y(epist), the epistatic effect of the interaction of the X and Y chromosomes; AUT(add) (or AUT(dom))_P, the epigenetic effect of the paternal genotype on the additive or dominant effect of the autosomes; Y (or X)→AUT(rec.epist), the epistatic effect of the Y (or X) chromosome on the recessive autosomal alleles; Y (or X)→AUT(dom.epist), the epistatic effect of the Y (or X) chromosome on the dominant autosomal alleles; and X_P, the epigenetic effect of the paternal genotype on the effect of the X chromosome. A plus sign indicates a combination of the respective effects. The effect of the paternal genotype alone on the traits under study was not considered because epigenetic effects were expected only for a combination of this factor with the offspring genotype. The genetic factors were treated as categorical variables; their significance was assessed by permutational ANOVA (PermANOVA). A paired permutation test was employed in post-hoc comparisons, and the Bonferroni correction was used for multiple comparisons. The maximal number of iterations was 100 000; the significance level was 0.05.

Several effects were combined in one variable because the study was not designed to examine all possible combinations of the factors and because several vectors determining the *A*_*n*_ indicator variables were collinear. To combine several effects in one joint effect, the values of the corresponding vectors were summed. Backcrosses to restore the *D. virilis* genotype serve to evaluate the effect of the *D. virilis* chromosomes on the resulting phenotype. The indicator variables were therefore taken to be 1 for the *D. virilis* genotype and 0 for the *D. lummei* genotype when considering effects of the X and Y chromosomes and a dominant effect of the autosomes and to be 1 for homozygotes for the *D. virilis* chromosomes, 0 for heterozygotes, and –1 for homozygotes for the *D. lummei* chromosomes when considering additive effects of the autosomes. To allow for possible spermatogenesis defects and subsequent epigenetic marking of the chromosomes in interspecific hybrids, the indicator variables for the factor Paternal Genotype were taken to be 1 for homozygous males of the parental species and 0 for heterozygous F_1_ males. The resulting matrix of genotype-dependent vector values is shown in Table 2.

**Table 2.**
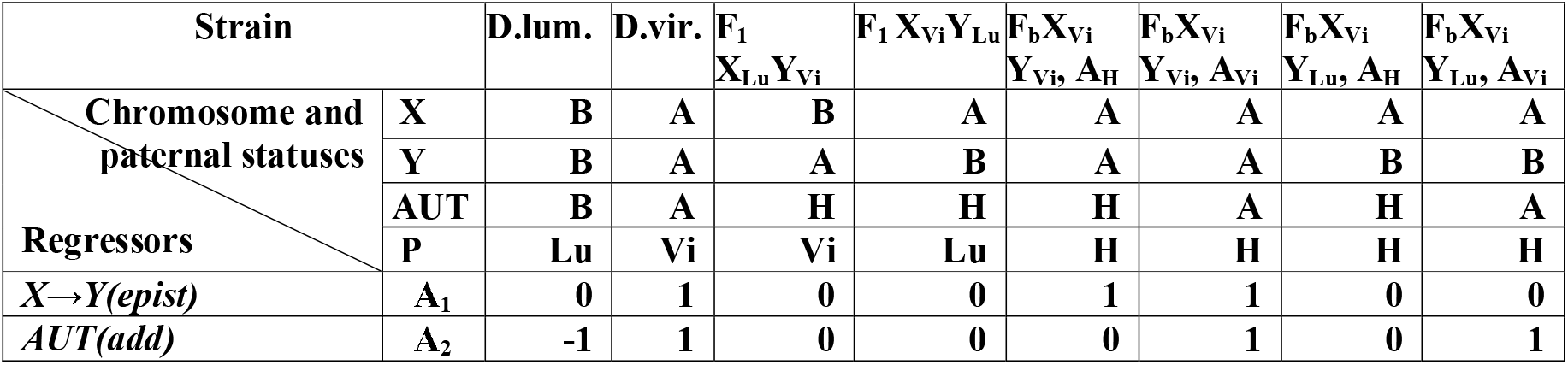
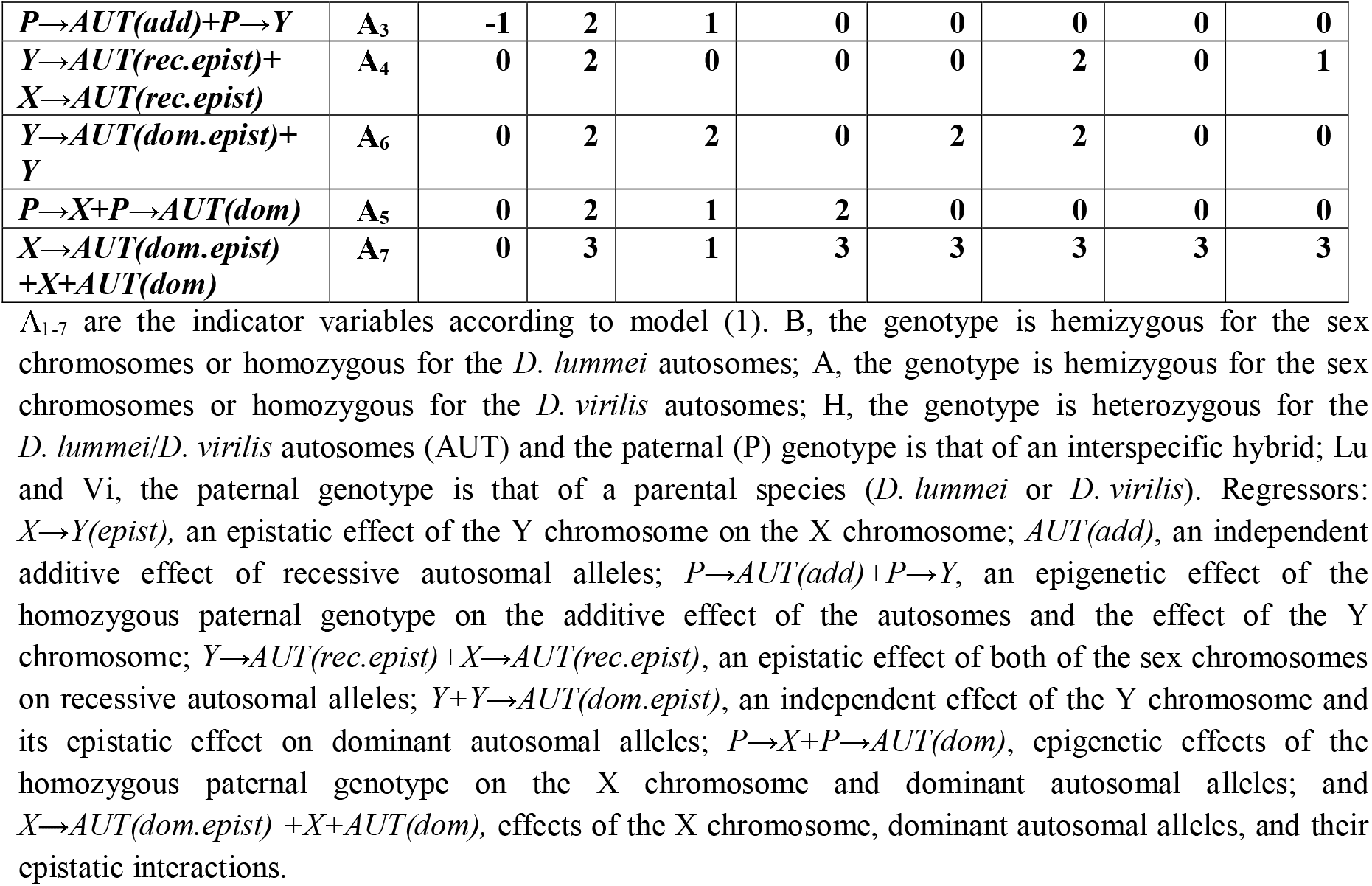
Indicator variables used to obtain the coefficients of regression between hereditary factors or their combinations and trait values.

A factor analysis was used to capture a covariance of traits; factors were extracted by the maximum likelihood method; factor loadings were examined by rotating principal component axes via the Promax procedure with the Kaiser normalization; estimates were obtained by regression analysis. The number of factors to be extracted isolated was determined using a scree plot of the eigenvalues of the trait correlation matrix.

Because a distinct elbow was not observed in the plot, we chose the factors whose eigenvalues were above a simulation curve obtained by a parallel analysis with 1000 permutations. The results of the factor analysis and their associations with genotypes were visualized by redundancy analysis (RDA), using the aggregate matrix of mean estimates for each factor. A principal component analysis (PCA) was performed for the initial MPs. The first two components were used in visualization.

## RESULTS

The variation in the shape of a morphological structure forms during development as various external and internal factors affect the differentiated cell population in an embryonic anlage of the respective organ. Parts of the anlage are topologically close together and experience similar effects, suggesting a similarity in a variation of the traits characterizing the parts of the morphological structure. To study the structure of interactions among the traits, and the latent factors responsible for a correlated variation, total variance over all samples was examined by a factor analysis. Seven factors were taken based on the scree plot and together accounted for 64.4% of the total variance (Table 3). Characteristics of the variation in the traits under study are shown in supplementary materials (Suppl. Tab.1).

**Table 3.**
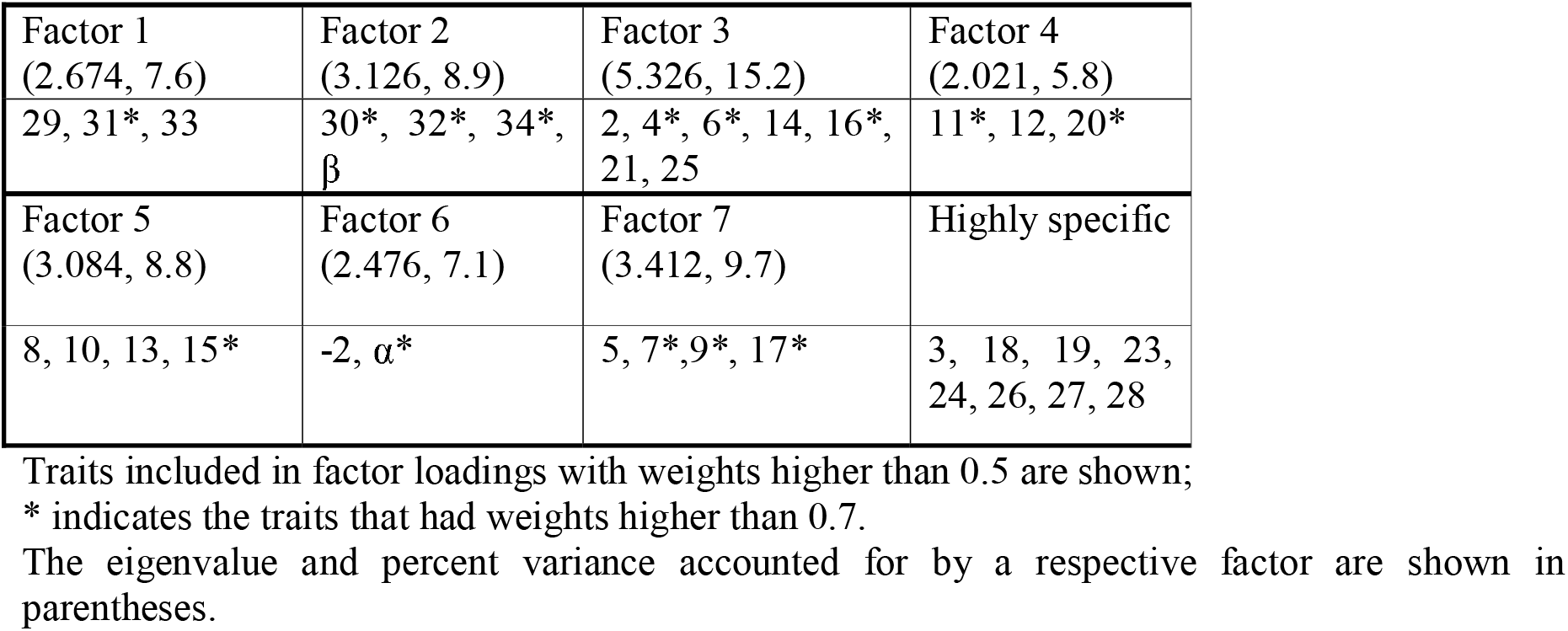
Compositions and eigenvalues of factor loadings as revealed by the maximum likelihood method.

Based on the traits with the greatest factor loadings, the factors were classified as follows.

Factor 1 characterized the variation in the width of the apodeme outline;

Factor 2 characterized the variation in the traits related to the apodeme bend;

Factor 3 included the most distinct species-specific traits and characterized the variation in (a) the shape of the ventral bend in the distal part of the aedeagus outline, (b) the position of the dorsal bend of the aedeagus outline relative to the central point, and (c) paramere lengths;

Factor 4 characterized the variation in traits related to the ventral bend in the proximal part of the aedeagus;

Factor 5 characterized the variation in traits related to the height of the dorsal bend of the aedeagus outline;

Factor 6 characterized the variation in the cook angle and the apical part of the aedeagus outline; and

Factor 7 characterized the variation in traits related to the dorsal bend in the distal part of the aedeagus outline and the lowermost point of the ventral bend in the same aedeagus part. IMP 3, 18, 19, 23, 24, 26, 27, and 28 were highly specific (Table 3, Highly specific).

RDA was carried out to relate the set of the variables that determine the factor structure values to the variables that characterize the genotype. In the analysis, the ordination of the genotypes and factors was performed in the space defined by correlations between factor weights, which were weighted sums of trait weights, and linear combinations of genotype scores, which were obtained for the genotypes characterizing the respective point in trait distribution. The center of gravity of the genotype distribution was brought into coincidence with the centers of gravity of the distribution of the factor structures. As is seen from Fig. 2, a two-dimensional space is defined by the orthogonal factor pair ML3–ML4 (X) and ML6, ML5–ML2, ML7 (Y). The genotypes showed the following arrangement in the two-dimensional factor space: +X: AAAH, ABAH, +Y(−X): BBBL, AAHH, −Y(−X): ABHL, BAHV (genotype abbreviations are as in Table 2). Genotypes ABHH and AAAV occur in the region of medium values. The angle between vectors on the plot and between genotype positions is proportional to the correlation between them. Therefore, the variation in traits of backcross males homozygous for the autosomes (with the *D. virilis* X chromosome) positively correlates with Factor 3 and negatively correlates with Factor 4. The finding indicates that the Y chromosome insignificantly affects the traits that determine the structures of the two factors. The contribution of the Y chromosome to the respective traits is somewhat greater in the backcross genotypes heterozygous for the autosomes but is still incomparable with the contributions that the Y chromosome makes to the traits in the other genotypes. Positive correlations with Factors 6 and 5 and negative correlations with Factors 2 and 7 were observed for traits of F_1_ males; opposite correlations, for traits of *D. lummei* males and backcross males with the *D. virilis* sex chromosomes. It is clear that combinations of the sex chromosomes in genotypes heterozygous or homozygous for the *D. lummei* autosomes and the male parent identity play a crucial role in the traits involved in the respective factor structures.

**Fig. 2.**
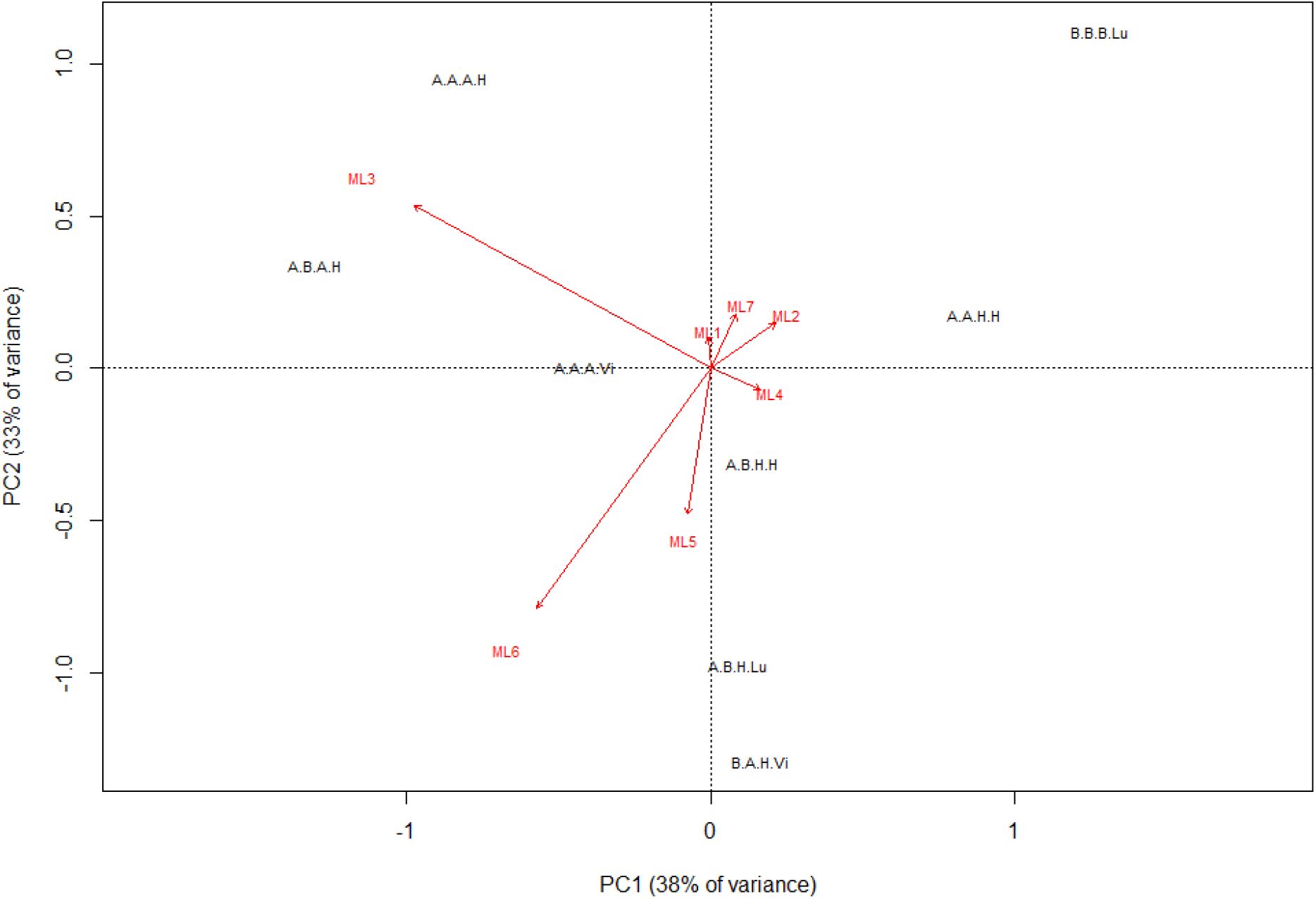
RDA: association of factor structure values and genotypes. ML1–7 are the variation vectors of Factors 1–7, which were obtained by the maximum likelihood method. The variables (traits) that constitute the X axis (in the order of decreasing factor loading) are: IMP6 (0.922), IMP4 (0.769), IMP16 (0.732), *IMP11* (*0.918*), *IMP20* (*0.729*). The variables (traits) that constitute the second Y axis are: IMP35 (0.890), IMP15 (0.832) *IMP32* (*1.039*), *IMP34* (*0.9*), *IMP30* (*0.843*), *IMP17* (*0.831*), *IMP9 (0.739*), *IMP7* (*0.718*). The least significant factors are in italics; factor loadings are shown in parentheses. The genotypes are abbreviated as in Table 2. The chromosomes and paternal genotype are indicated in the following order: X chromosome, Y chromosome, autosomes, male parent identity.

The results are supported by the distribution of the genotypes on a scatter plot (Suppl. Fig. 1) of the two first principal components, which are similar in loading structure to Factors 6 and 3. The parental genotypes are in opposite corners of the plot, having the lowest values in the case of *D. virilis* and the highest in the case of *D. lummei*. The cloud of *D. virilis* data overlaps the clouds of F_b_ males homozygous for autosomes regardless of the Y-chromosome status. The other genotypes are distributed between the *D. lummei* cloud and the cloud of *D. virilis* with homozygous F_b_ males; F_1_ males and F_b_ males heterozygous for autosomes differ in variation and form slightly overlapping clouds. Different genotypes differ in how close their variation parameters are to the parameters of the parental species at different principal component axes. Again, trait expression depends on the combination of the sex chromosome status, the autosome status, and the male parent identity as an epigenetic factor.

Two approaches were used to more precisely evaluate the effect that the sex chromosomes exert on trait expression in the shape of the male copulatory system. First, ANOVA was performed to compare the dominance of parental species-specific phenotypes in progenies from reciprocal crosses between *D. virilis* and *D. lummei* and backcrosses of F_1_ males with *D. virilis* females. All genotypes had the same autosome set and differed in sex chromosome combination and the identity of the male parent (the original parental species or the F_1_ hybrid). Second, ANOVA and post-hoc tests were performed using the genotype at the sex chromosomes, the genotype at the autosomes, and combinations of these factors, including the male parent identity, as independent variables.

The phenotypes of males obtained in direct and reciprocal crosses and backcross males heterozygous for all autosomes were compared with the phenotypes of males of the parental species. The results are summarized in Table 4. The logic of estimating the degree of dominance for a trait has been described previously (Kulikov et al., 2013). Based on the distribution of the hybrid and parental genotypes over groups isolated by post-hoc comparisons, it is possible to identify the following variants: incomplete dominance, the dominance of the *D. virilis* or *D. lummei* phenotype, and lack of difference among all phenotypes. The resulting data clustering variants were ranged by the degree of phenotype dominance. All cases where a genotype in question appeared together with the parental genotypes in one group were considered to suggest no significant difference (ns) in evaluating the degree of dominance. The variants *f_1(b)_<l<v, v<l<f_1(b)_, l,f_1(b)_<v, v<f_1(b)_,l, v*≤*f_1(b)_,l*, and *l,f_1(b)_*≤*v* suggested dominance of the *D. lummei* phenotype (D_Lu_); the variants *l<f_1(b)_<v, l*≤*f_1(b)_*≤*v*, *v*≤*f_1(b)_*≤ *l*, and *v<f_1(b)_<l*, intermediate dominance (ID); and *f_1(b)_ <v<l, l<v< f_1(b)_, f_1(b)_,v <l, l<f_1(b)_*,*v*, *l*≤*v,f_1(b)_*, and *f_1(b)_,v*≤*l*, dominance of the *D. virilis* phenotype (D_Vi_). The Gabriel test and Tukey’s HSD test yielded similar results; only those of the Gabriel test are shown.

**Table 4.**
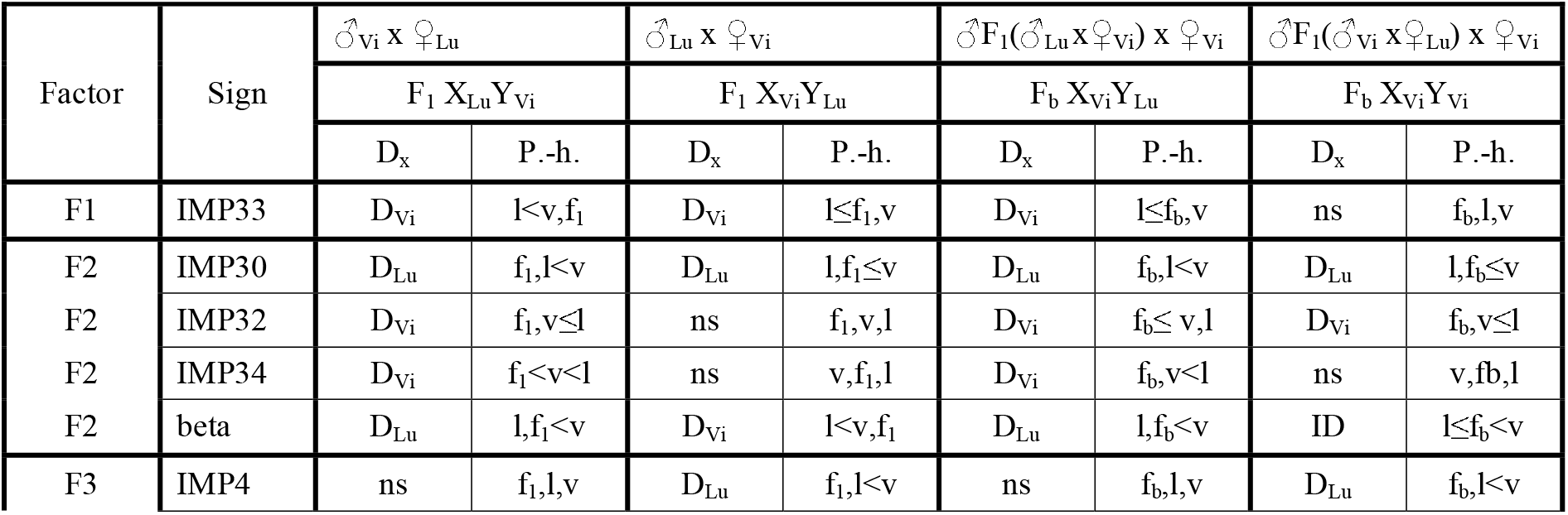
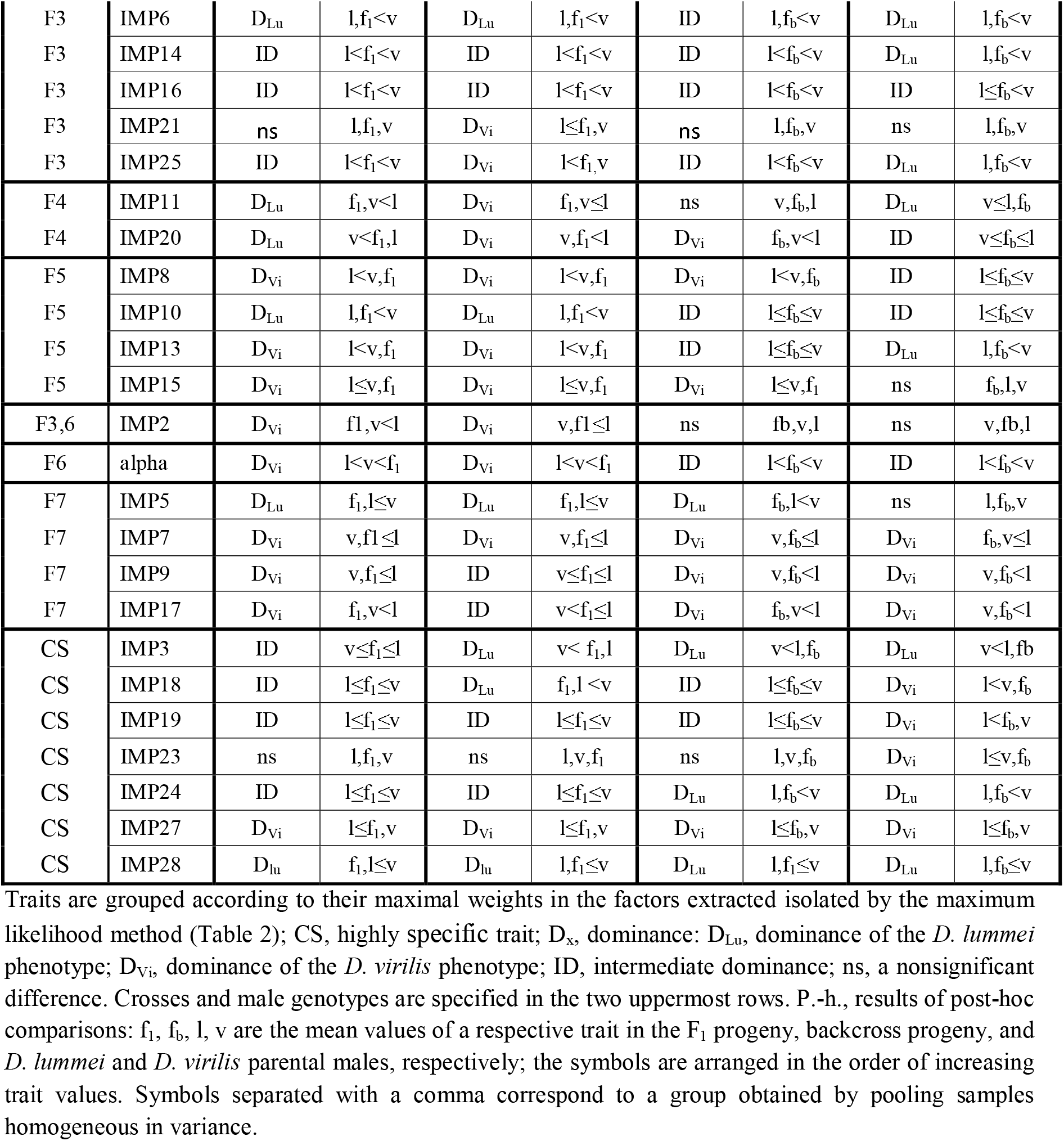
Dominance of copulative system shape-related traits as dependent on the sex chromosome composition in *D. virilis/D. lummei* hybrid males, heterozygous for the autosomes

IMPs 12, 22, 26, 29, and 31 did not significantly depend on the genotype and are not included in Table 4. The IMPs that showed significant correlations were grouped; the results of this grouping are considered below together with factor analysis results. Here we describe the most general results of the analysis of variance. First, traits at which the *D. virilis* phenotype dominated (43) prevailed over traits with dominance of the *D. lummei* phenotype (32) and traits with intermediate dominance (26) in all four samples. Dominance of the same phenotype regardless of the sex chromosome combination was observed for only 5 (IMPs 7, 16, 27, 28, and 30) out of the 30 IMPs included in the analysis, suggesting a substantial role of the autosome combination and the identity of the male parent in trait expression. The effect of the male parent identity was evaluated by comparing F_1_ males with genotype X_Vi_Y_Lu_ and backcross males with the same genotype, given that crossing over is absent in males and the males in question were entirely identical in chromosome composition. Differences in parental phenotype dominance were observed for 16 out of the 30 traits, and the dominance character changed to the opposite one in the case of apodeme declination angle. A substitution of the *D. lummei* Y chromosome for its *D. virilis* counterpart in backcross males similarly changed the dominance character in 16 traits. Of these, ten traits, which mostly loaded on Factors 1, 3, 4, and 5, showed a change to dominance of the opposite species-specific phenotype relative to the species origin of the Y chromosome. Phenotypic comparisons of F_1_ progenies showed that simultaneous substitution of the sex chromosomes changed the character of dominance at 12 traits, of which three (IMP 11, IMP20, and β) changed their dominance status to the opposite one, according to the species origin of the X chromosome. Other changes were less distinct and included firstly, intermediate dominance was shifted toward dominance of one of the parental species. Secondly, the phenotype was changed relative to that of the parental species so that a homogeneous variance group within one of the parental species and offspring with a particular genotype was replaced with a group that included offspring with an alternative genotype and both of the parental species (Table 4, category “ns”). It is of interest to note that the greatest difference in the total number of traits showing a dominance of one of the parental phenotype was observed in backcross flies heterozygous for the autosomes. Thus F_b_ X_Vi_Y_Vi_ males had eight traits at which the *D. virilis* phenotype dominated and ten traits at which *D. lummei* phenotype dominated, while F_b_ X_Vi_Y_Lu_ males displayed an opposite pattern and had ten and five such traits, respectively.

As expected, the number of traits at which the *D. virilis* phenotype dominated increased in backcross males homozygous for the *D. virilis* autosomes (Suppl. Tab.2); the set included 19 traits in F_b_ X_Vi_Y_Vi_ males and 23 traits in F_b_ X_Vi_Y_Lu_ males. A substitution of the Y chromosome changed the character of dominance at nine traits, of which seven again demonstrated a negative effect of the species identity of the Y chromosome on the phenotype.

Factorial MANOVA was carried out to evaluate the effects of the sex chromosomes, autosomes, paternal genotype, and their interactions on the phenotypic traits. Significant effects were observed for each of the four factors taken alone and the interaction of the Y chromosome and autosomes (Suppl. Табл. 3).

To study the effect on particular phenotypic traits for each of the factors, MANOVA was performed with forced incorporation of all four predictors and the effect of the ChrY*Aut interaction. The results are summarized in Suppl. Table 4. The identities of the X and Y chromosomes, the male parent identity, the autosome status, and the interaction of the autosome with the Y chromosome were included as independent variables in the analysis. The majority of the traits showed an association with the identities of the X chromosome and autosomes at a significance level p < 0.05; half of the traits were affected by the Y chromosome and male parent identities; and one-third of the traits, by the interaction of the Y chromosome and autosomes. A cooperative effect of these variables was mostly observed for the groups of traits extracted by the factor analysis. An analysis of the factor values confirmed the effect in the cases where the majority of the traits determining the respective factor structure significantly depended on the given variable. Note that a dependence on the hereditary factors was not confirmed for the apodeme width-related traits (F1). The apodeme declination and curvature (F2) depend on the identities of the Y chromosome and male parent. Distinct species-specific traits of the aedeagus and parameres (F3) depend on the X chromosome, autosomes, and male parent identity. The shape of the ventral bend in the proximal part of the aedeagus outline (F4) is determined by the interaction between the Y chromosome and autosomes. Traits related to the height of the dorsal bend in the aedeagus outline (F5) and the shape of the cook and the outline bend over the cook (F6) showed a similar dependence on the Y chromosome, autosomes, and male parent identity; an additional dependence on the X chromosome is specific to F5-related traits. Traits describing the shape of the distal part of the outline (F7) depend on the autosomes, male parent identity, and their interaction.

The effect of the male parent identity is possibly mediated through the epigenetic marking of chromosomes in interspecific hybrids during gametogenesis and a subsequent effect of the resulting signatures on the ontogeny of offspring. As mentioned above, an independent effect of the Y chromosome is expected to be insignificant based on the composition and functional activity of its coding sequences. It is possible to assume that interactions of the two factors play a crucial role in the phenotype. To detail the strength and direction of the effects exerted by the hereditary factors and their combinations on the dominance of the *D. virilis* phenotype, post-hoc tests were used to evaluate the probability for groups homogeneous in trait variance to form according to a published model (Hobert, 2008) (Table 6; Suppl., Table 5). The results obtained for each particular trait are provided in Supplementary; data on latent and highly specific traits are summarized in Table 6.

**Table 6.** Paired permutation test with the Bonferroni correction to evaluate the optimal genotype partitioning into groups homogeneous in latent variables and highly specific variables (up to 100 000 iterations in each case)

| F# | | $X \rightarrow Y$<br>(epist) | AUT<br>(add) | $P \rightarrow AUT(ad$<br>$d) + P \rightarrow Y$ | $Y + Y \rightarrow A$<br>$UT(dom.$<br>$epist)$ | $P \rightarrow AUT(add)$<br>$+ P \rightarrow Y$ | $Y \rightarrow AUT$<br>(rec.epist)<br>$+ X \rightarrow AUT$<br>(rec.epist) | $X \rightarrow AUT$<br>(dom.epist)<br>$+ X + AUT$<br>(dom) |
| --- | --- | --- | --- | --- | --- | --- | --- | --- |
| F1 | Prob. | 0.493 | 0.476 | 0.473 | 0.136 | 0.302 | 0.638 | 0.304 |
|  | Dom. | n.s. | n.s. | n.s. | n.s. | n.s. | n.s. | n.s. |
| F2 | Prob. | 0.113 | 0.207 | **0.000** | **0.038** | **0.000** | 0.057 | **0.000** |
| | Dom. | n.s. | n.s. | + $sD^{Lu}_{X-chr}$ | + | +* ($X-chr$) | n.s. | +* ($sD^{Vi}_{X-chr}$) |
| F3 | Prob. | **0.001** | **0.000** | **0.000** | 0.285 | **0.000** | **0.000** | **0.000** |
| | Dom. | + | + $D^{Lu}$ | +* ($sD^{Vi}_{Aut}$) | n.s. | + $P_Y, D^{Lu}_{Aut}$ | +* $sD^{Vi}_{X-chr}$ | +* $sD^{Lu}_{X-chr}$ |
| F4 | Prob. | 0.697 | 0.489 | **0.000** | **0.000** | **0.000** | 0.738 | **0.000** |
| | Dom. | n.s. | n.s. | +* | + | + $P_Y$ | n.s. | +* $sD^{Lu}_{X-chr}$ |
| F5 | Prob. | **0.000** | **0.000** | 0.043 | **0.025** | **0.001** | **0.000** | 0.051 |
| | Dom. | + | +* | n.s. | + | +* $D^{Lu}_{Aut}$ | + ID | n.s. |
| F6 | Prob. | 0.478 | **0.000** | **0.000** | **0.000** | **0.000** | 0.857 | **0.000** |
| | Dom. | n.s. | + $D^{Vi}$ | + $D^{Vi}_{Aut}$ | + | + $D^{Vi}_{Aut} (X-chr)$ | n.s. | + ID |
| F7 | Prob. | 0.542 | **0.000** | 0.045 | **0.031** | **0.002** | 0.028 | **0.001** |
|  | Dom. | n.s. | + sD<sup>Vi</sup> | n.s. | + | +* D<sup>Vi</sup><sub>Aut</sub> | +* | + D<sup>Vi</sup><sub>Aut</sub> |
| FDR | 0.05 | 0.021 | 0.043 | 0.029 | 0.043 | 0.036 | 0.036 | 0.021 |
|  | 0.01 | 0.004 | 0.009 | 0.006 | 0.006 | 0.007 | 0.004 | 0.004 |
| Imp 3 | Prob. | **0.000** | **0.000** | 0.811 | 0.009 | **0.000** | **0.000** | 0.123 |
|  | Dom. | + | + D<sup>Lu</sup> | n.s. | + | + D<sup>Lu</sup><sub>Aut</sub> | + D<sup>Vi</sup><sub>X-chr</sub> | n.s. |
| Imp 18 | Prob. | **0.000** | **0.000** | 0.222 | 0.008 | **0.009** | **0.000** | 0.085 |
|  | Dom. | + | + D<sup>Lu</sup> | n.s. | + | + P<sub>Aut</sub> | + D<sup>Vi</sup><sub>X-chr</sub> | n.s. |
| Imp 19 | Prob. | **0.000** | **0.000** | 0.302 | **0.000** | **0.002** | **0.000** | **0.009** |
|  | Dom. | + | + ID | n.s. | + | + D<sup>Vi</sup><sub>Aut</sub> | + ID | + D<sup>Vi</sup><sub>Aut</sub> |
| Imp 23 | Prob. | 0.385 | **0.004** | 0.033 | 0.412 | **0.002** | **0.000** | **0.002** |
|  | Dom. | n.s. | + D<sup>Vi</sup> | n.s. | n.s. | +* D<sup>Vi</sup><sub>Aut</sub> | +* | + ID |
| Imp 24 | Prob. | **0.010** | **0.004** | 0.000 | **0.000** | **0.004** | **0.003** | **0.016** |
|  | Dom. | + | + D<sup>Vi</sup> | + D<sup>Vi</sup><sub>Aut</sub> | + | + D<sup>Vi</sup><sub>Aut</sub> | + ID | +* sD<sup>Vi</sup><sub>Aut</sub> |
| Imp 26 | Prob. | 0.055 | **0.000** | 0.239 | 1.000 | 0.145 | **0.000** | 0.120 |
|  | Dom. | n.s. | + D<sup>Lu</sup> | n.s. | n.s. | n.s. | + D<sup>Vi</sup><sub>X-chr</sub> | n.s. |
| Imp 27 | Prob. | 0.306 | **0.000** | 0.054 | 0.253 | **0.004** | **0.000** | **0.002** |
|  | Dom. | n.s. | + D<sup>Lu</sup> | n.s. | n.s. | +* X-chr | +* sD<sup>Vi</sup><sub>X-chr</sub> | + D<sup>Lu</sup><sub>X-chr</sub> |
| Imp 28 | Prob. | **0.020** | 0.538 | **0.000** | 1.000 | **0.002** | 0.042 | 0.067 |
|  | Dom. | + | n.s. | + D<sup>Lu</sup><sub>X-chr</sub> | n.s. | + D<sup>Lu</sup><sub>Aut</sub> | n.s. | n.s. |

The model implies that each hereditary factor exerts a discrete effect on the traits in question, depending on the genotype. The expected effect of each factor was analyzed using the indicator variables listed in Table 2. We expected that the variation in each trait is possible to describe using several categories corresponding to the number of indicator variables for a given hereditary factor.

Factor **X**→**Y(epist)** has two indicator variable categories, the effect being present or absent. A significant difference in these categories between samples confirms the significant effect of the interaction between the sex chromosomes. The effect of the conspecific *D. virilis* sex chromosomes is examined versus the effects of all other combinations in this case.

Factor **Y+Y**→**AUT(dom.epist)** has two similar categories. The independent effect of the Y chromosome alone on the traits is close to zero. Therefore, a significant difference in the categories between samples confirms the significant effect of the interaction between the Y chromosome and autosomes. The effect of the *D. virilis* Y chromosome and autosomes is examined versus the effects of all other combinations in this case.

With all subsequent factors, a significant difference between the extreme variants confirms that the respective factor significantly affects the trait in question. A significant difference between intermediate and other genotype groups in the absence of differences between the extreme groups indicates that the effect in question depends nonlinearly on the genotype.

Factor **AUT(add)** has three indicator variable categories: two opposite effects of the *D. virilis* and *D. lummei* homozygous autosomes and an intermediate effect of the heterozygous autosomes. An intermediate effect suggests a decrease in additive interactions of divergent genes, and the lack of significant difference from one of the homozygous genotypes indicates that dominant alleles mostly contribute to the additive effect.

Factor **Y**→**AUT(rec.epist)+X**→**AUT(rec.epist)** has three categories: an effect of interactions between the homozygous *D. virilis* autosomes and the *D. virilis* sex chromosomes, a partial effect of interactions between the homozygous *D. virilis* autosomes and the X chromosome, and all variants with the heterozygous autosomes or homozygous *D. lummei* autosomes. A significant difference between the extreme variants indicates that the sex chromosomes interact with recessive autosomal genes, leading to species-specific distinctions. Lack of a significant difference from one of the extreme variants suggests a predominant contribution of epistatic interactions with one of the sex chromosomes, depending on the genotype combination (dominance of the *D. virilis* phenotype suggests a leading role for the X chromosome; dominance of the *D. lummei* phenotype, for the *D. virilis* Y chromosome, which is absent in 1.9.30 males, of the *D. lummei* Y chromosome present in the given genotype).

Factor **P**→**X+P**→**AUT(dom)** has three categories: the minimal and maximal values correspond, respectively, to the absence or presence of interactions of the *D. virilis* X chromosome and autosomes with the homozygous paternal genotype, and an intermediate value corresponds to interactions of dominant genes of the *D. virilis* autosomes with the *D. virilis* homozygous paternal genotype. A grouping of genotypes having intermediate values with those having one of the extreme values maximizes the role of the autosomes or the X chromosome, depending on the group composition (dominance of the *D. virilis* phenotype suggests a maximal role for the autosomes; dominance of the *D. lummei* phenotype, for the *D. virilis* X chromosome, which is absent in (P_Vi_)F_1_ X_Lu_Y_Vi_A_Vi_A_Lu_ males).

Factor ***X***→***AUT(dom.epist)+X+AUT(dom)*** has three categories. The minimal and maximal values correspond to expression of the *D. virilis* phenotype due to the effects of dominant autosomal genes and the *D. virilis* X chromosome and their epistatic interactions. The intermediate value corresponds to a sole effect of dominant autosomal genes. Dominance of the *D. virilis* phenotype (a grouping of genotype X^Lu^/Y^Vi^_Aut^Vi^/Aut^Lu^ with genotypes X^Vi^/Y^*^_Aut^Vi^/Aut^*^) suggests a predominant effect for *D. virilis* dominant autosomal genes; dominance of the *D. lummei* phenotype, for the *D. lummei* X chromosome, *D. lummei* autosomes, or their combination.

Factor **P**→**AUT(add)+P**→**Y** has four indicator variable categories. The extreme values define the effects that the paternal genotype exerts on the expression of the *D. virilis* phenotype under the influence of recessive autosomal alleles and the Y chromosome. The two alternative extreme values accordingly belong to the genotypes of the *D. virilis* and *D. lummei* parental strains. The intermediate values of indicator variables are defined by the epigenetic effect that the *D. virilis* male parent identity exerts exlusively on the *D. virilis* Y chromosome (genotype (P_Vi_)F_1_ X_Lu_Y_Vi_A_Vi_A_Lu_, the value is 1) and by lack of this effect on the genotypes that are heterozygous for the autosomes and have the Y chromosome originating from a heterozygous male or a *D. lummei* male (genotypes (P_Lu_)F_1_ X_Vi_Y_Lu_A_Vi_A_Lu_, (P_Vi/Lu_)F_b_ X_Vi_Y_Vi_A_Vi_A_Lu_, (P_Vi/Lu_)F_b_ X_Vi_Y_Vi_A_Vi_A_Vi_, (P_Vi/Lu_)F_b_ X_Vi_Y_Lu_A_Vi_A_Lu_, and (P_Vi/Lu_)F_b_ X_Vi_Y_Lu_A_Vi_A_Vi_; the value is 0). Lack of a significant epigenetic effect on the autosomes will lead to the clustering of the *D. lummei* genotype with genotype (P_Vi_)F_1_ X_Lu_Y_Vi_A_Vi_A_Lu_ and the *D. virilis* genotype with genotypes (P_Lu_)F_1_ X_Vi_Y_Lu_A_Vi_A_Lu_, (P_Vi/Lu_)F_b_ X_Vi_Y_Vi_A_Vi_A_Lu_, (P_Vi/Lu_)F_b_ X_Vi_Y_Vi_A_Vi_A_Vi_, (P_Vi/Lu_)F_b_ X_Vi_Y_Lu_A_Vi_A_Lu_, and (P_Vi/Lu_)F_b_ X_Vi_Y_Lu_A_Vi_A_Vi_. If a significant epigenetic or genetic effects of the Y chromosome is lacking, the genotype (P_Vi_)F_1_ X_Lu_Y_Vi_A_Vi_A_Lu_ will cluster with genotypes (P_Lu_)F_1_ X_Vi_Y_Lu_A_Vi_A_Lu_, (P_Vi/Lu_)F_b_ X_Vi_Y_Vi_A_Vi_A_Lu_, (P_Vi/Lu_)F_b_ X_Vi_Y_Vi_A_Vi_A_Vi_, (P_Vi/Lu_)F_b_ X_Vi_Y_Lu_A_Vi_A_Lu_, and (P_Vi/Lu_)F_b_ X_Vi_Y_Lu_A_Vi_A_Vi_. If the autosomes exert a dominant effect characteristic of *D. virilis* or *D. lummei* in this case, strains with the intermediate values of indicator variables will cluster together with the respective parental genotype. The incomplete design of crosses makes the results difficult to interpret. For example, male genotype (P_Vi_)F_1_ X_Lu_Y_Vi_A_Vi_A_Lu_ was the only genotype that had the intermediate value 1 for the indicator variables ***A_3_*** (factor ***P***→***AUT(add)+P***→***Y***), ***A_6_*** (factor ***P***→***X+P***→***AUT(dom)***), and ***A_7_*** (factor ***X***→***AUT(dom.epist) +X+AUT(dom)***). It is clear that the phenotypic features of this genotype are determined by combined effects of the three factors, and its clustering with other genotypes may therefore be distorted in the case of factor ***P***→***AUT(add)+P***→***Y***.

Table 6 and Suppl. Table 3 summarize the results of analyzing how the primary and latent traits varied under the influence of the hereditary factors. A dependence on the hereditary factors was observed for the majority of the latent traits and the corresponding primary traits with a high commonality. Trait F1, which includes traits related to apodeme width, was an exception; i.e., its dependence on any of the hereditary factors was not confirmed.

The simplest explanation of the results is possible for the hereditary factors of epistatic interactions of the X and Y chromosomes with the X chromosome and the X chromosome with the autosomes, each having only two categories of indicator variables. A significant effect of the first factor ***X***→***Y(epist)*** was demonstrated for latent traits F3 and F5 and confirmed for at least half of the corresponding primary traits and IMPs 3, 18, 19, 24, and 28, which are highly specific. The factor ***Y***→***AUT (dom. epist)+Y*** significantly affect latent traits F2 and F4–F7, as well as highly specific primary IMPs 3, 18, 19, and 24. A significant effect of this interaction was not observed for the primary traits incorporated in F2, F5, and F7 with high weights. The result indicates that the proportion of variation included in correlated traits is codirectional in character and is detectable in the respective latent traits. It is safe to say that the epistatic effects exerted by the conspecific sex chromosomes and by the interaction between the Y chromosome and dominant alleles of the D. virilis autosomes positively affect expression of the D. virilis phenotype in the corresponding morphological structures.

A significant influence of the additive effect of recessive autosomal alleles was demonstrated for latent traits F3 and F5–F7 and confirmed for the majority of the corresponding primary traits and highly specific IMPs 3, 18, 19, 24, 26, and 27. Traits of the genotypes that were heterozygous for the autosomes and had an intermediate value of the indicator variable displayed differentiation by the dominance of one of the parental phenotypes. The finding indicates that a dominant component acts as part of this hereditary factor. Latent trait F3, which characterizes the most distinct species-specific traits, and highly specific IMPs 3, 18, 26, and 27 showed dominance of the *D. lummei* phenotype in heterozygotes for the autosomes. Primary traits included in F3 with high weights similarly displayed dominance of the *D. lummei* phenotype (IMPs 4 and 21) or intermediate dominance (IMPs 6, 14, 16, and 25) in genotypes heterozygous for the autosomes. Latent traits F6 and F7, which characterize different parts of the aedeagus outline, and highly specific IMPs 23 and 24 displayed dominance of the *D. virilis* phenotype in these genotypes. Although one trait showing dominance of the *D. lummei* phenotype was found in each of the correlated variability groups characterizing latent traits F6 and F7, the traits apparently made only a minor contribution to the total variance. Primary traits included in latent trait F5 with high weights variously showed dominance for both of the parental genotypes, and the total variance of F5 nonlinearly depended on the autosome status, separating the heterozygous genotypes and either homozygous genotype. The additive effect of the autosomes significantly influences dominance of the *D. virilis* phenotype and, in our variation analysis model, incorporates a dominant component that determines the expression of one of the parental phenotypes, depending on the trait in question.

The epistatic effects of the sex chromosomes on recessive autosomal alleles (factor *Y*→*AUT(rec.epist)+X*→*AUT(rec.epist)*) significantly influence latent traits F3, F5, and F7 and highly specific primary IMPs 3, 18, 19, 23, 24, 26, and 27. Dominance of the *D. virilis* phenotype was mostly observed for traits of backcross males, which were homozygous for the *D. virilis* autosomes, carried the *D. virilis* X chromosome and *D. lummei* Y chromosome, and had the intermediate value of the indicator variable. The finding suggests a significant effect for the *D. virilis* X chromosome. The effect is fully confirmed by traits incorporated in latent trait F3 with high weights. The variables that constitute latent factors F5 and F7 included both traits dependent on and those independent of the epistatic interaction between the sex chromosomes and autosomes. The dependent traits showed opposite dominance variants for the genotype with an intermediate value of the indicator variable. An analysis of the pooled variance of these traits as components of latent traits showed incomplete dominance of the genotype with the intermediate value of the indicator variable in the case of F5 and a significant difference of this genotype from the total set of other genotypes in the case of F7. It is possible to assume that the latent traits experience epistatic effects from both of the sex chromosomes in the former case and a nonlinear effect from the combined influence of the above epistatic effects in the latter case. It should be noted that the primary traits that constitute latent traits F4 and F6 also showed a dependence on the epistatic effects that the sex chromosomes exert on the autosomes, but the dependence was not confirmed in the analysis of the pooled variance. The epistatic effects that the *D. virilis* sex chromosomes exert on recessive alleles of the conspecific autosomes significantly influence the dominance of the *D. virilis* phenotype, the X chromosome presumably playing a leading role.

A combined effect of dominant autosomal alleles, the X chromosome, and their epistatic interactions (factor *X*→*AUT(dom.epist)+X+AUT(dom)*) was confirmed for the majority of the primary traits having a high commonality; latent traits F2–F4, F6, and F7; and highly specific IMPs 19, 23, 24, and 27. However, the effect differed among the traits, thus determining a heterogeneous group composition for the majority of the latent traits. Latent trait F4 was an exception. This trait and its component primary traits of the highest weights showed a dominance of the *D. lummei* phenotype in males with the intermediate value of the indicator variable (genotype F_1_ X_Lu_Y_Vi_), suggesting a predominant effect for the *D. lummei* X chromosome and autosomes. The variation of latent trait F3 displayed a similar dependence on the *D. lummei* X chromosome and/or autosomes. In contrast, a significant effect of this factor was observed for all primary traits involved in latent trait F5, but the variation of F5 itself did not show a dependence on the factor in question because opposite effects were combined. Traits related to the apodeme bend and declination (F2) nonlinearly depended on the factor, and the genotypes showed a partitioning into two homogeneous variance groups with the intermediate and extreme values of the indicator variable. However, the finding that the mean values gradually decreased in the order X_Lu_Y_Lu_ Aut_Lu_/Aut_Lu_ → X_Vi_Y* Aut_Vi_/Aut* → X_Lu_Y_Vi_ Aut_Lu_/Aut_Vi_ suggests the possibility of superdominance of the *D. virilis* phenotype in the intermediate genotype and a leading role of dominant alleles of the *D. virilis* autosomes. A similar effect of *D. virilis* dominant autosomal alleles was confirmed for latent trait F7 and highly specific IMPs 19 and 24 at high significance. Latent trait F6 and highly specific IMP 23 displayed incomplete *D. virilis* phenotype dominance of genotype X_Lu_Y_Vi_ Aut_Lu_/Aut_Vi_, corresponding to the expected effect of factor *X*→*AUT (dom.epist)+X+AUT(dom)*. Dominant autosomal alleles, the X chromosome, and their interactions certainly affect the phenotype, although their particular effects are impossible to differentiate in this case.

Influence of the male parent identity on the effect of the X chromosome and dominant autosomal alleles (factor ***P***→***X+P***→***AUT(dom)***) was demonstrated for latent traits F2, F3, F4, and F6 and highly specific primary IMPs 24 and 28. The expected effect of this factor on all genotypes was confirmed for latent trait F4 and both of the primary traits (IMPs 11 and 20) that determine its variation. Latent trait F3 nonlinearly depended on the epigenetic effect that the male parent identity exerts on the X chromosome and autosomes, differentiating the genotypes into two groups with the extreme and intermediate values of the indicator variable. The dominance of the *D. lummei* phenotype was observed in the genotype having the intermediate value of the indicator variable for latent trait F2; primary IMPs 2, 30, 32, and beta, which determine the F2 variation; and highly specific primary IMP 28. The finding suggests a predominant epigenetic effect on the X chromosome for these traits. An effect on the autosomes was confirmed by the dominance of the *D. virilis* phenotype in these genotypes and was observed for latent trait F6 and highly specific IMP 24. A significant effect of this factor was not demonstrated for latent traits F1, F5, and F7, while individual primary traits incorporated in the latent factor displayed, in the intermediate genotype, the dominance of the *D. virilis* phenotype in the case of F1 and F5 and the *D. lummei* phenotype in the case of F7. With F7, the mean values gradually increased in the order of genotypes with indicator variable values 1–0–2, suggesting superdominance of the *D. lummei* phenotype and a substantial role of epigenetic modification of the X chromosome.

Thus, the estimates confirm that epistatic interactions of the sex chromosomes and autosomes and epigenetic effects of the male parent origin from interspecific crosses influence the expression of species-specific traits in the shape of the male copulatory system.

## DISCUSSION

How do traits determining the shape of the male copulatory system become a target of selection? As mentioned in the Introduction, the efficiency of female insemination in insects depends on how well the female and male genitalia match each other (Jagadeeshan, Singh. 2006). Incomplete insemination must make repeated mating more likely, lead to displacement of sperm from the previous mating, and facilitate efficient selection against the incomplete insemination-associated genotype in *Drosophila*. Hoikkala and colleagues (Mazzi et al., 2009) have analyzed the within-population variation of mating duration in *D. montana* and associated the duration of the first mating with female resistance to repeated mating. We have shown that sensory microchaetae on the ventral surface of the aedeagus mediate the association between evolutionarily significant parts of the *Drosophila* copulatory system with mating behavior (Kulikov et al., 2013).

Both mating behavior and the shape of the copulatory system are not directly involved in adaptation but are maintained close to the adaptive optimum of a population as a result of apparent stabilizing selection (Johnson, Barton 2005; Zhang, Hill 2005). In other words, the selection at these characters acts through adaptively valuable traits characteristic of a particular group of individuals. This selection takes the form of sexual selection because the characters are expressed specifically as predictors of sexual reproduction efficiency. When a new adaptation forms and the adaptive norm changes rapidly, selection changes to directional or disruptive, but still remains indirect. It is, therefore, possible to expect that the variation observed experimentally will correspond to the variation in quantitative traits affected by one of the above selection types.

The variation maintained by stabilizing selection is equally well described by two models, a joint-effect model and an infinitesimal model. In the former, the variation is associated with effects of moderate-frequency alleles, which are nearly neutral in terms of fitness and substantially influence the trait in question. In the latter, the variation involves many genes that each exert a minor effect on fitness and act additively. Modeling of ample experimental data has shown that well reproducible results are obtained with both of the models, none of them is possible to prefer over the other (Zhang, Hill, 2005). An analysis of QTLs for commercially valuable traits in farm animals supports well the conclusion that recessive variation maintained in a population plays an important role (Andersson, Georges, 2004; Hill, 2014; Zan et al., 2017). The majority of the traits are related to morphological and physiological characteristics affected by stabilizing selection. The facts that a variety of farm animal breeds formed rapidly on an evolutionary scale, that homologous haplotypes are responsible for similar traits in different breeds and are present in populations of founder species, and that epistatic interactions are observed between QTLs indicate that diversity at quantitative traits is maintained because neutral variation is preserved in populations.

Lower estimates could be obtained for genetic variation at traits under stabilizing selection when the effects of suppressing epistasis are underestimated, the fact being masked by overestimating the effect of stabilizing selection and/or underestimating the magnitude of variation due to mutation (Mackay, 2010, 2015). Moreover, opposite epistatic effects may occur between tightly linked genes in a QTL (Dworkin et al., 2003) to reduce their total effect observed, and opposite effects occurring between different QTLs substantially increase the total variance.

The formation of evolutionary new and usually adaptively significant traits has another genetic dynamics. Fisher (Fisher, 1930) has noted that adaptation is not selection, meaning fitness-based selection that eliminates the least fit individuals. Adaptation is characterized by the effect of positive selection, which facilitates the progress of a population to an optimal phenotype. Based on Maynard Smith’s model of adaptive walk in sequence space (M.Smith, 1970) and Gillespie’s theory of fixation of rare beneficial alleles (Gillespie, 1983, 1991), it is thought now that adaptations arise in bursts alternating with periods of slow evolution (Gillespie, 1989). Adaptation is associated with a few effects, and selection rapidly decreases in strength with each subsequent step; i.e., an exponential distribution is characteristic of the selection coefficients of consecutively fixed mutations (Orr, 2002). An allele that expands the ecological niche for the species will be fixed rapidly in a new QTL. Experiments on QTL mapping support the conclusions based on model analyses (Terekhanova et al., 2014; Lin et al., 2016; Lendenmann et al., 2016; Bay et al., 2017). It is safe to say that, along with additive effects, dominant interactions of alleles gain principal importance. A detailed analysis of the genetic architecture and ontogenetic mechanisms of species-specific traits in two *Labeotropheus* species from the Malawi Lake have made it possible to evaluate the role that regulatory, or epistatic, interactions of genes involved in the Tgfβ signaling pathway play in the formation of foraging adaptations (Conith et al., 2018).

Additive, dominant, epistatic, and epigenetic components of variation underlie the phenotypic differences in our experiment. The experiment was not designed to exactly determine the variance components. However, given that the homozygous or heterozygous autosome sets were identical in the males examined, it is clear that changes in the dominance of the parental phenotypes are associated with epigenetic effects of the male parent identity and epistatic effects of the X and Y chromosomes. Neither primary nor latent traits remained phenotypically the same in genotypes with different combinations of the sex chromosomes and different male parent identities, while the autosome set was identical.

The effects of sex chromosomes and autosomes were specific to different groups of traits. For example, the additive effect of recessive autosomal alleles determined the traits of the aedeagus (apart from those characterizing the ventral bend in the proximal part of its outline) and parameres. A combined effect of dominant autosomal alleles, the X chromosome, and its dominant epistatic interaction with the autosomes was observed for traits of the aedeagus and parameres and the bend and declination of the apodeme. It is of interest that the two factors exerted similar effects in genotypes with intermediate values of indicator variables. This was dominance of the *D. lummei* phenotype in latent trait F3, which combined the most distinct species-specific traits; certain primary traits that were incorporated in latent traits F4 and F5 and similarly showed species specificity (Kulikov et al., 2004); and paramere width (IMP 27). Dominance of the *D. virilis* phenotype was characteristic of latent trait F7; several primary traits incorporated in latent traits F5 and F6, and paramere width in the ventral bend region (IMP 24). The examples show that the dominant component of the autosomes contributed to the effect of additive interactions and that its contribution depended on the epistatic interaction with the X chromosome. Following Huang and Mackay (Huang, Mackay, 2016), the components of variance are impossible to strictly extracted isolate in the majority of cases, especially when the experimental design is not an orthogonal one, which at least formally ensures uncorrelated variance components. Our estimates are therefore total effects of several components of variance. Minimization of the autosomal effect in the case of dominance of the *D. lummei* phenotype and, oppositely, minimization of the effect of the X chromosome and its epistatic influence on dominant autosomal alleles show that the X chromosome plays alternative roles in the variation of different traits. Similar dominance changes observed for genotypes heterozygous for the autosomes when estimating the role of the additive component of variance confirms that the dominant component of the autosomes contributes to the species-specific variation.

The effect that epistatic interactions between the X and Y chromosomes exert on species-specific traits was confirmed for half of the primary traits and latent traits F3 and F5, which incorporate taxonomically significant traits. It should be noted here that unequivocal interpretation is impossible for the results indicating that the *D. virilis* phenotype is enhanced in males with conspecific sex chromosomes. Given that different variance components may contribute to this phenomenon, an independent effect can be expected for genes of the *D. virilis* X chromosome. For example, Carson and Lande (Carson, Lande, 1984) have shown that the formation of an evolutionarily new secondary sex characteristic (an additional row of bristles on the male tibia) is determined to the extent of 30% by sex-linked genes in a natural *Drosophila silvestris* race. As for the traits that contribute to isolating barriers, the most detailed data are available for sterility-related traits. The examples below illustrate interspecific hybrid male sterility as a model of interactions between the autosomes and sex chromosomes. A cluster of *Stellate* sequences on the *D. melanogaster* X chromosome codes for a homolog of the β subunit of protein kinase CK2, and its overexpression causes male sterility. RNA interference mediated by Y-linked *Su*(*Ste*) repeats prevents *Stellate* overexpression (Kotelnikov et al.,2009; Olenkina et al., 2012), demonstrating that male fertility strongly depends on the interaction between the X and Y chromosomes. A similar model of fertility regulation utilizes the Odysseus-site homeobox protein (OdsH), which is encoded by an X-chromosomal locus in species of the *melanogaster* group. OdsH binds to heterochromatin sequences of the Y chromosome. In many cases, the presence of heterospecific X and Y chromosomes leads to decondensation of Y-chromosomal heterochromatin, dramatically distorts the expression of autosomal genes, and causes sterility (Bayes, Malik, 2009; Lu et al., 2010). The genetic system also illustrates the interaction between the X and Y chromosomes.

The effect of epistatic interactions between the sex chromosomes and recessive alleles of the autosomes was confirmed for the majority of latent and primary traits. A leading role of epistatic interactions of the X chromosome is evident from the observation that the *D. virilis* phenotype dominated at the majority of traits in males with genotype X^Vi^/Y^Lu^ Aut^Vi^/Aut^Vi^. A substantial role of epistatic interactions between the X chromosome and autosomes has similarly been demonstrated for another trait involved in prezygotic barriers, namely, a species-specific male courtship song pattern in *Drosophila* species of the *montana* phylad (Päällysaho et al., 2003). There is evidence that the sex chromosomes exert a significant regulatory effect on expression activity of autosomal genes. Expression of X-chromosomal genes in hemizygous *D. simulans* males depends on trans regulations to a substantial extent (Wayne et al., 2004), and trans effects and joint action of cis and trans effects of the X chromosome on genome-wide expression activity are the second most important to trans-regulatory effects of chromosome III in *D. melanogaster* males (Wang et al., 2008).

The important role that the Y chromosome plays in regulating the phallus shape-related traits in interspecific hybrids is possible to directly associate with trans-regulatory activity. The Y chromosome harbors only 23 single-copy protein-coding genes, and 13 of them are strongly restricted to the testis in expression and are mostly associated with hybrid male sterility (Carvalho et al., 2001, Piergentili, Mencarelli, 2008, Vibranovski et al., 2008, Francisco, Lemos, 2014). Additive variation of the ten other genes should make a vanishingly small contribution to the phenotype at quantitative traits of morphological structures. However, epistatic interactions of Y-chromosomal sequences with the X chromosome and autosomes have received experimental support. For example, experiments were performed to evaluate the activity of the male-specific lethal proteins/roX1,2 RNA complex, which is responsible for dosage compensation of X-linked genes in *Drosophila* males. The Y chromosome proved to affect the viability in *roX1, roX2* mutant males, the effect depending on the Y-chromosome source (Menon, Meller, 2009). The viability was low in males the paternal Y chromosome and high in males with the maternal Y chromosome. The result indicates that the Y chromosome modifies dosage compensation through *roX1, roX2-*mediated modification of heterochromatin (Deng et al., 2009) and/or recognition of the X chrome by the entire male-specific lethal proteins/roX1,2 RNA complex. The results support our finding of a substantial paternal effect, which acts independently in the majority of traits or in combination with the X-chromosome effect in some other cases. Finally, Lemos and colleagues (Francisco, Lemos 2014; Lemos, Araripe, Hartl, 2008) have directly demonstrated the regulatory activity of the Y chromosome in a study of differential genome expression activity under the Y-chromosome influence.

Traits of the apodeme, which is a muscle attachment site and internal part of the copulatory system, are the least associated together by evolutionary variation and chromosome effects. It is noteworthy that chromosome effects are mediated by the epigenetic effect of male parent identity in the case of the apodeme traits. Note that significant epigenetic effects were similarly observed for all other groups of correlated traits. The epigenetic effect may be related to the mechanism of meiotic sex chromosome inactivation (MSCI) or, in a broader sense, meiotic silencing of unpaired chromatin/DNA (MSUC/MSUD) (Turner, 2007; Tao et al., 2007), which inactivates the chromosome regions with altered meiotic synapsis in leptotene. The MSCI mechanism and its substantial evolutionary role in organizing the chromosomes and facilitating postzygotic isolation has been the matter of extensive discussion (Turner, 2007; Tao et al., 2007; Meiklejohn, Tao, 2010; Vibranovski, 2014). MSCI is typically compensated for by the higher transcriptional activity of X-linked genes in *Drosophila* because potent promoters are overrepresented in the X chromosome (Deng et al., 2011; Landeen et al., 2016). The formation of heterochromatin blocks is often associated with sterility in hybrids from interspecific crosses and death of hybrid progenies; defects in mitotic chromosome segregation are observed hybrids during their embryo development (Turner, 2007; Ferree, Barbash, 2009; Bayes, Malik, 2009). Interspecific differences cannot accumulate at once and may be seen as changes in gene expression activity and alter the trait magnitudes in early species divergence. Epigenetic signatures are preserved in chromosomes at postmeiotic stages and may be inherited through generations (Somer, Thummel, 2014; Francisco, Lemos, 2014). Although the regulation of genome-wide epigenetic states is often associated with the Y chromosome (Friberg et al., 2012; Branco et al., 2013; Francisco, Lemos, 2014), a formal role of the interspecific hybrid status is noteworthy in our case, leading to distorted synapsis of divergent chromosomes and causing the formation and further preservation of noncanonical heterochromatin regions with altered expression activity.

Thus, it is possible to say that the two sex chromosomes act unidirectionally to shift phenotype dominance in the conspecific direction. The effect of the Y chromosome is mediated to a substantial effect by its epistatic interactions with the X chromosome and autosomes.

