## Supplemental Tables 1-5 for "The Role of the Sex Chromosomes in the Inheritance of Species Specific Traits of the Shape of the Copulatory Organ in Drosophila"

Suppl. Tabl.1. Variation of the morphometric traits in the shape of the male copulatory system

| Signs | 0.0.4 (10) | | 1.0.8 (31) | | 1.0.9 (25) | | 2,8,1 (5) | | 2,8,13 (5) | | 2,9,1 (28) | | 2,9,30 (10) | | vir160 (13) | |
| --- | --- | --- | --- | --- | --- | --- | --- | --- | --- | --- | --- | --- | --- | --- | --- | --- |
|  | M | σ | M | σ | M | σ | M | σ | M | σ | M | σ | M | σ | M | σ |
| IMP2^3,6^ | 1.023 | 0.013 | 0.998 | 0.018 | 1.008 | 0.010 | 1.021 | 0.013 | 1.058 | 0.047 | 1.019 | 0.020 | 1.028 | 0.012 | 1.019 | 0.010 |
| IMP3 | 0.110 | 0.012 | 0.098 | 0.017 | 0.106 | 0.017 | 0.123 | 0.017 | 0.066 | 0.010 | 0.100 | 0.017 | 0.086 | 0.022 | 0.076 | 0.020 |
| IMP4^3^ | 0.151 | 0.017 | 0.153 | 0.019 | 0.150 | 0.018 | 0.145 | 0.025 | 0.188 | 0.025 | 0.159 | 0.023 | 0.191 | 0.016 | 0.175 | 0.021 |
| IMP5^7^ | 0.023 | 0.012 | 0.018 | 0.013 | 0.020 | 0.013 | 0.025 | 0.014 | 0.017 | 0.006 | 0.009 | 0.007 | 0.024 | 0.008 | 0.025 | 0.007 |
| IMP6^3^ | 0.180 | 0.020 | 0.201 | 0.030 | 0.203 | 0.021 | 0.190 | 0.022 | 0.328 | 0.061 | 0.218 | 0.041 | 0.298 | 0.042 | 0.259 | 0.041 |
| IMP7^7^ | 0.066 | 0.011 | 0.040 | 0.024 | 0.050 | 0.008 | 0.050 | 0.020 | 0.034 | 0.026 | 0.038 | 0.013 | 0.064 | 0.040 | 0.052 | 0.024 |
| IMP8^5^ | 0.287 | 0.019 | 0.372 | 0.044 | 0.381 | 0.034 | 0.329 | 0.028 | 0.381 | 0.034 | 0.387 | 0.046 | 0.381 | 0.037 | 0.368 | 0.023 |
| IMP9^7^ | 0.057 | 0.019 | 0.030 | 0.033 | 0.047 | 0.018 | 0.013 | 0.009 | 0.008 | 0.020 | 0.018 | 0.026 | 0.022 | 0.029 | 0.011 | 0.029 |
| IMP10^5^ | 0.352 | 0.044 | 0.350 | 0.062 | 0.346 | 0.050 | 0.286 | 0.046 | 0.215 | 0.034 | 0.298 | 0.046 | 0.234 | 0.047 | 0.230 | 0.026 |
| IMP11^4^ | 0.087 | 0.021 | 0.113 | 0.036 | 0.054 | 0.029 | 0.088 | 0.015 | 0.042 | 0.023 | 0.061 | 0.031 | 0.058 | 0.046 | 0.061 | 0.020 |
| IMP12^4^ | 0.145 | 0.031 | 0.124 | 0.029 | 0.128 | 0.026 | 0.106 | 0.014 | 0.119 | 0.019 | 0.120 | 0.022 | 0.173 | 0.171 | 0.115 | 0.030 |
| IMP13^5^ | 0.397 | 0.037 | 0.431 | 0.032 | 0.428 | 0.032 | 0.366 | 0.020 | 0.402 | 0.034 | 0.400 | 0.039 | 0.413 | 0.027 | 0.417 | 0.043 |
| IMP14^3^ | 0.273 | 0.051 | 0.356 | 0.057 | 0.359 | 0.048 | 0.306 | 0.058 | 0.505 | 0.016 | 0.353 | 0.056 | 0.477 | 0.056 | 0.472 | 0.046 |
| IMP15^5^ | 0.372 | 0.035 | 0.405 | 0.030 | 0.405 | 0.030 | 0.360 | 0.014 | 0.383 | 0.034 | 0.402 | 0.035 | 0.419 | 0.047 | 0.380 | 0.028 |
| IMP16^3^ | 0.362 | 0.037 | 0.447 | 0.070 | 0.467 | 0.051 | 0.475 | 0.077 | 0.689 | 0.054 | 0.503 | 0.068 | 0.633 | 0.035 | 0.591 | 0.033 |
| IMP17^7^ | 0.084 | 0.018 | 0.045 | 0.014 | 0.063 | 0.011 | 0.058 | 0.015 | 0.036 | 0.023 | 0.046 | 0.020 | 0.054 | 0.014 | 0.054 | 0.024 |
| IMP18 | 0.624 | 0.070 | 0.657 | 0.084 | 0.619 | 0.053 | 0.745 | 0.050 | 0.747 | 0.160 | 0.669 | 0.057 | 0.717 | 0.071 | 0.732 | 0.071 |
| IMP19 | 0.176 | 0.020 | 0.229 | 0.041 | 0.194 | 0.028 | 0.236 | 0.026 | 0.289 | 0.040 | 0.204 | 0.042 | 0.241 | 0.061 | 0.245 | 0.054 |
| IMP20^4^ | 0.107 | 0.019 | 0.105 | 0.026 | 0.062 | 0.018 | 0.092 | 0.016 | 0.045 | 0.023 | 0.073 | 0.028 | 0.060 | 0.030 | 0.067 | 0.021 |
| IMP21^3^ | 0.073 | 0.006 | 0.077 | 0.012 | 0.080 | 0.015 | 0.076 | 0.018 | 0.082 | 0.004 | 0.076 | 0.011 | 0.098 | 0.010 | 0.085 | 0.009 |
| IMP22 | 0.025 | 0.011 | 0.032 | 0.020 | 0.029 | 0.014 | 0.027 | 0.010 | 0.023 | 0.010 | 0.024 | 0.025 | 0.017 | 0.011 | 0.026 | 0.015 |
| IMP23 | 0.171 | 0.016 | 0.191 | 0.037 | 0.195 | 0.023 | 0.236 | 0.025 | 0.208 | 0.030 | 0.205 | 0.030 | 0.240 | 0.034 | 0.192 | 0.028 |
| IMP24 | 0.169 | 0.033 | 0.240 | 0.057 | 0.239 | 0.061 | 0.187 | 0.036 | 0.308 | 0.086 | 0.184 | 0.058 | 0.206 | 0.073 | 0.259 | 0.069 |
| IMP25^3^ | 0.163 | 0.029 | 0.247 | 0.052 | 0.294 | 0.033 | 0.218 | 0.036 | 0.392 | 0.022 | 0.251 | 0.049 | 0.356 | 0.051 | 0.344 | 0.052 |
| IMP26 | 0.069 | 0.015 | 0.074 | 0.029 | 0.071 | 0.026 | 0.078 | 0.023 | 0.097 | 0.014 | 0.080 | 0.020 | 0.103 | 0.031 | 0.089 | 0.015 |
| IMP27 | 0.162 | 0.041 | 0.171 | 0.022 | 0.187 | 0.020 | 0.184 | 0.014 | 0.198 | 0.025 | 0.176 | 0.024 | 0.209 | 0.037 | 0.181 | 0.021 |
| IMP28 | 0.606 | 0.075 | 0.581 | 0.103 | 0.655 | 0.064 | 0.638 | 0.021 | 0.571 | 0.065 | 0.593 | 0.074 | 0.578 | 0.108 | 0.695 | 0.115 |
| IMP29^1^ | 0.068 | 0.021 | 0.063 | 0.014 | 0.059 | 0.010 | 0.053 | 0.009 | 0.063 | 0.006 | 0.062 | 0.011 | 0.056 | 0.012 | 0.059 | 0.008 |
| IMP30^2^ | 0.049 | 0.021 | 0.039 | 0.024 | 0.078 | 0.029 | 0.053 | 0.021 | 0.057 | 0.029 | 0.042 | 0.017 | 0.046 | 0.009 | 0.081 | 0.046 |
| IMP31^1^ | 0.048 | 0.006 | 0.052 | 0.014 | 0.050 | 0.008 | 0.043 | 0.002 | 0.045 | 0.005 | 0.046 | 0.007 | 0.047 | 0.004 | 0.049 | 0.007 |
| IMP32^2^ | 0.103 | 0.033 | 0.052 | 0.032 | 0.109 | 0.043 | 0.082 | 0.030 | 0.087 | 0.036 | 0.063 | 0.026 | 0.051 | 0.016 | 0.088 | 0.044 |
| IMP33^1^ | 0.037 | 0.006 | 0.052 | 0.018 | 0.046 | 0.004 | 0.035 | 0.004 | 0.044 | 0.002 | 0.043 | 0.008 | 0.050 | 0.003 | 0.047 | 0.009 |
| IMP34^2^ | 0.103 | 0.038 | 0.034 | 0.030 | 0.080 | 0.035 | 0.070 | 0.033 | 0.073 | 0.024 | 0.048 | 0.028 | 0.030 | 0.022 | 0.068 | 0.031 |
| alpha^6^ | 0.353 | 0.158 | 1.070 | 0.089 | 1.127 | 0.140 | 0.688 | 0.151 | 0.911 | 0.154 | 0.817 | 0.249 | 1.059 | 0.113 | 1.018 | 0.059 |
| beta^2^ | 0.194 | 0.112 | 0.250 | 0.101 | 0.371 | 0.115 | 0.293 | 0.061 | 0.286 | 0.133 | 0.239 | 0.114 | 0.239 | 0.128 | 0.434 | 0.115 |

The sample size is indicated in parentheses. M, mean; σ, standard deviation. Superscripts in the first column indicate that the trait is incorporated with a high weight in the respective factor structure (Table 3).

Suppl. Pic.1. Genotype distribution in the space of the first two principal components.

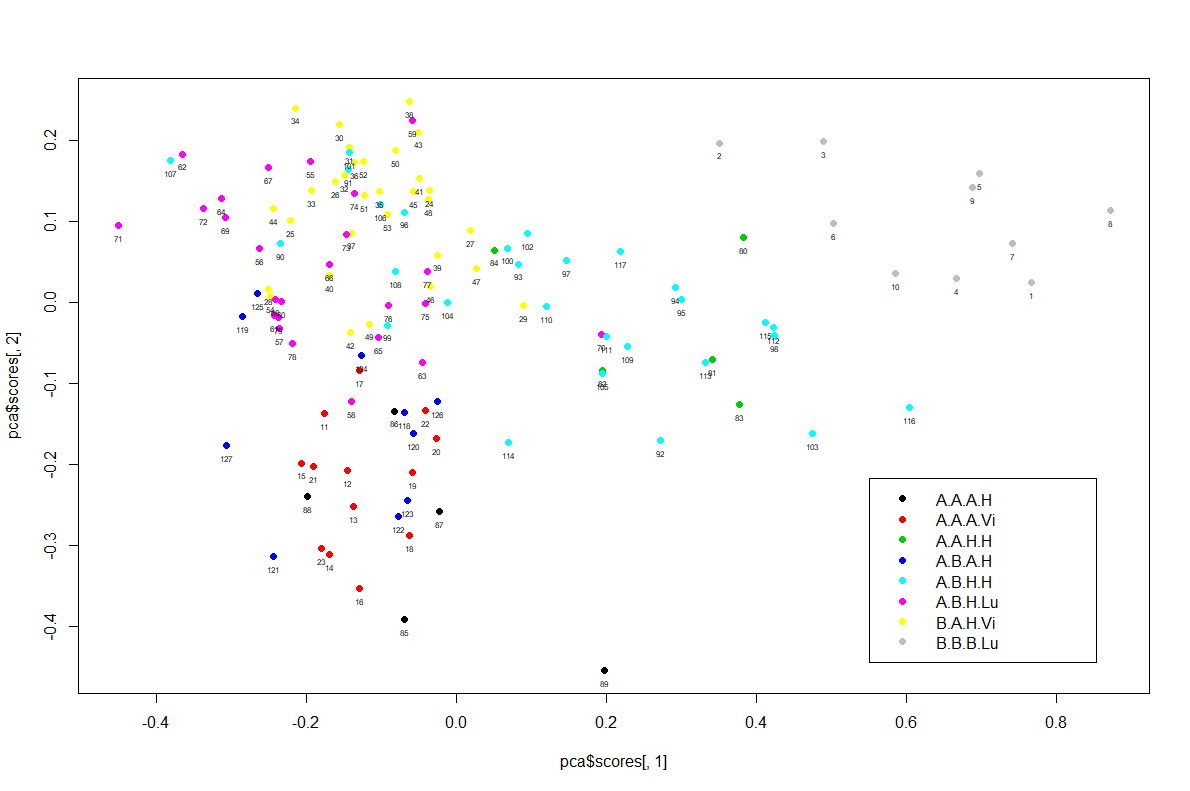

The genotypes are abbreviated as in Table 2. The chromosomes and paternal genotype are indicated in the following order: X chromosome, Y chromosome, autosomes, male parent identity.

Suppl. Tab.2. Dominance at traits of the copulatory system shape as dependent on the sex chromosome composition in *D. virilis/D. lummei* hybrid males homozygous for the *D. virilis* autosomes

| Factor | Sign | ♂F_1_(♂_Lu_ x♀_Vi_) x ♀_Vi_ | | ♂F_1_(♂_Vi_ x♀_Lu_) x ♀_Vi_ | |
| --- | --- | --- | --- | --- | --- |
|  |  | F_b_ X_Vi_Y_Lu_ **2.9.30** | | F_b_ X_Vi_Y_Vi_ **2.8.13** | |
|  |  | D_x_ | P.-h. | D_x_ | P.-h. |
| F1 | imp33 | D_Vi_ | l≤v,f_b_ | ns | l,v,f_b_ |
| F2 | imp30 | D_Lu_ | f_b_,l<v | D_Lu_ | l,f_b_≤v |
| F2 | imp32 | D_Vi_ | f_b_,v<l | ns | f_b_,v,l |
| F2 | imp34 | D_Vi_ | f_b_,v<l | D_Vi_ | v,f_b_≤l |
| F2 | beta | D_Lu_ | l,f_b_<v | D_Lu_ | l,f_b_<v |
| F3 | imp4 | D_Vi_ | l≤v,f_b_ | D_Vi_ | l≤v,f_b_ |
| F3 | imp6 | D_Vi_ | l<f_b_<v | D_Vi_ | l<f_b_<v |
| F3 | imp14 | D_Vi_ | l<v,f_b_ | D_Vi_ | l<v,f_b_ |
| F3 | imp16 | D_Vi_ | l<v,f_b_ | D_Vi_ | l<v<f_b_ |
| F3 | imp21 | D_Vi_ | l<v,f_b_ | ns | l,fb,v |
| F3 | imp25 | D_Vi_ | l<v,f_b_ | D_Vi_ | l<v,f_b_ |
| F4 | imp11 | D_Vi_ | f_b_,v≤l | D_Vi_ | f_b_,v<l |
| F4 | imp20 | D_Vi_ | f_b_,v<l | D_Vi_ | f_b_,v<l |
| F5 | imp8 | D_Vi_ | l<v,f_b_ | D_Vi_ | l<v,f_b_ |
| F5 | imp10 | D_Vi_ | v,f_b_<l | D_Vi_ | f_b_,v<l |
| F5 | imp13 | ns | l,f_b_,v | D_Lu_ | l,f_b_≤v |
| F5 | imp15 | D_Vi_ | l≤v,f_b_ | D_Vi_ | l≤v,f_b_ |
| F3,6 | imp2 | ns | v,l,f_b_ | D_Lu_ | v≤l,f_b_ |
| F6 | alpha | D_Vi_ | l<v,f_b_ | D_Vi_ | l<f_b_,v |
| F7 | imp5 | ns | l,f_b_,v | ns | l,f_b_,v |
| F7 | imp7 | D_Vi_ | f_b_,v≤l | ns | v,f_b_,l |
| F7 | imp9 | D_Vi_ | f_b_,v≤l | D_Vi_ | f_b_,v<l |
| F7 | imp17 | D_Vi_ | f_b_,v<l | D_Vi_ | f_b_,v<l |
| HH | imp3 | D_Vi_ | v,f_b_<l | D_Vi_ | f_b_,v<l |
| HH | imp18 | D_Vi_ | l<f_b_,v | D_Vi_ | l<v,f_b_ |
| HH | imp19 | D_Vi_ | l<f_b_,v | D_Vi_ | l<v,f_b_ |
| HH | imp23 | D_Vi_ | l≤v,f_b_ | ns | l,v,f_b_ |
| HH | imp24 | ID | l≤f_b_≤v | D_Vi_ | l<v,f_b_ |
| HH | imp27 | D_Vi_ | l<v,f_b_ | D_Vi_ | l<v,f_b_ |
| HH | imp28 | D_Lu_ | l,f_b_≤v | ns | l,f_b_,v |

Suppl. Tabl 3. Significant effects and interactions as revealed by factorial MANOVA

|  | ChrX | ChrY | Aut | ♂P | ChrY*Aut |
| --- | --- | --- | --- | --- | --- |
| F | 12.06 | 10.51 | 12.97 | 3.64 | 2.32 |
| p | 0 | 0 | 0 | 0 | 0.0009 |

All four factors and their interactions were used as predictors; the 35 phenotypic traits, as independent variables.

Suppl. Tab 4. Effects of the sex chromosomes, autosomes, and parental genotypes on trait expression

| Factor | Sign | ChrX | | ChrY | | Aut | | ♂P | | ChrY*Aut | |
| --- | --- | --- | --- | --- | --- | --- | --- | --- | --- | --- | --- |
|  |  | F | p | F | p | F | p | F | p | F | p |
| F1 | imp33 | 3.38 | 0.0684 | 2.11 | 0.1489 | 9.32 | **0.0002** | 1.11 | 0.3327 | 0.13 | 0.7165 |
|  | F1 | 0.19 | 0.663 | 0.06 | 0.813 | 0.76 | 0.468 | 0.26 | 0.774 | 0.14 | 0.711 |
| F2 | imp30 | 14.23 | **0.0002** | 1.03 | 0.312 | 2.01 | 0.1386 | 15.86 | **7.86E-07** | 0 | 0.9538 |
|  | imp32 | 7.14 | **0.0086** | 2.66 | 0.1056 | 10.12 | **8.76E-05** | 13.68 | **4.48E-06** | 0.47 | 0.4959 |
|  | imp34 | 2.87 | 0.0928 | 4.61 | **0.0337** | 21.32 | **1.22E-08** | 8.35 | **0.0004** | 0.87 | 0.3514 |
|  | beta | 13.08 | **0.0004** | 10.3 | **0.0017** | 0.17 | 0.8408 | 15.15 | **1.38E-06** | 0.01 | 0.9343 |
|  | F2 | 1.73 | 0.191 | 4.99 | **0.027** | .78 | 0.459 | 9.98 | **0.000** | 0.60 | 0.439 |
| F3 | imp4 | 7.96 | **0.0056** | 4.11 | **0.0449** | 19.96 | **3.34E-08** | 2.11 | 0.12 | 0.53 | 0.4672 |
|  | imp6 | 34.28 | **4.31E-08** | 18.93 | **2.88E-05** | 41.39 | **2.27E-14** | 6.1 | **0.003** | 5.35 | **0.0224** |
|  | imp14 | 33.58 | **5.72E-08** | 46.78 | **3.65E-10** | 43.02 | **8.71E-15** | 0.32 | 0.7266 | 3.83 | 0.0528 |
|  | imp16 | 87.76 | **5.87E-16** | 45.67 | **5.48E-10** | 38.5 | **1.27E-13** | 5.73 | **0.0042** | 3.89 | 0.0508 |
|  | imp25 | 66.22 | **4.45E-13** | 36.34 | **1.91E-08** | 33.61 | **2.68E-12** | 7.93 | **0.0006** | 4.39 | **0.0383** |
|  | imp21 | 6.55 | **0.0117** | 0.76 | 0.3863 | 12.6 | **1.09E-05** | 0.34 | 0.7145 | 3.53 | 0.0626 |
|  | F3 | 5.94 | **0.016** | 2.87 | 0.093 | 62.44 | **0.000** | 6.42 | **0.002** | 0.44 | 0.506 |
| F4 | imp11 | 63.21 | **1.19E-12** | 3.17 | 0.0776 | 1.92 | 0.1498 | 0.68 | 0.507 | 3.34 | 0.0700 |
|  | imp12 | 0.07 | 0.7932 | 3.04 | 0.0839 | 2.88 | 0.0597 | 0.36 | 0.7002 | 1.05 | 0.3065 |
|  | imp20 | 73.03 | **5.04E-14** | 0 | 0.9956 | 2.56 | 0.0819 | 2.25 | 0.1102 | 3.62 | 0.0596 |
|  | F4 | 0.007 | 0.933 | 0.11 | 0.742 | 0.30 | 0.745 | 0.78 | 0.461 | 5.18 | **0.025** |
| F5 | imp10 | 43.6 | **1.18E-09** | 22.41 | **6.12E-06** | 20.14 | **2.92E-08** | 6.62 | **0.0019** | 0.03 | 0.8557 |
|  | imp8 | 14.24 | **0.0002** | 2.26 | 0.135 | 20.91 | **1.65E-08** | 0.14 | 0.8669 | 4.38 | **0.0386** |
|  | imp13 | 3.34 | 0.0703 | 0.19 | 0.6662 | 5.34 | **0.006** | 6.76 | **0.0017** | 0.91 | 0.3417 |
|  | imp15 | 0.02 | 0.8986 | 2.3 | 0.1318 | 11.37 | **3.04E-05** | 0.07 | 0.9332 | 0.04 | 0.8434 |
|  | F5 | 4.15 | **0.044** | 8.59 | **0.004** | 14.59 | **0.000** | 3.38 | **0.037** | 0.23 | 0.630 |
| F3;6 | imp2 | 20.36 | **1.51E-05** | 0.05 | 0.8309 | 15 | **1.60E-06** | 9.01 | **0.0002** | 4.58 | **0.0343** |
| F6 | alpha | 5.28 | **0.0234** | 33.28 | **6.44E-08** | 69.6 | **< 2.2e-16** | 28.78 | **6.39E-11** | 0.03 | 0.8704 |
|  | F6 | 1.94 | 0.166 | 8.13 | **0.005** | 46.64 | **0.000** | 23.97 | **0.000** | 0.540 | 0.464 |
| F7 | imp5 | 0.85 | 0.3586 | 3.06 | 0.0828 | 5.44 | **0.0055** | 4.65 | **0.0113** | 8.97 | **0.0033** |
|  | imp7 | 0.02 | 0.9012 | 3.05 | 0.0831 | 7.14 | **0.0012** | 1.54 | 0.2195 | 8.14 | **0.0051** |
|  | imp17 | 0.18 | 0.6681 | 18.11 | **4.19E-05** | 11.93 | **1.89E-05** | 6.63 | **0.0019** | 5.87 | **0.0169** |
|  | imp9 | 5.66 | **0.0189** | 17.33 | **5.96E-05** | 0.81 | 0.4466 | 8.34 | **0.0004** | 0.21 | 0.6471 |
|  | F7 | 1.65 | 0.202 | 2.37 | 0.127 | 7.83 | **0.001** | 9.39 | **0.000** | 7.92 | **0.006** |
| HC | imp3 | 2.34 | 0.1283 | 18.54 | **3.43E-05** | 16.02 | **6.91е-07** | 0.54 | 0.5829 | 11.34 | **0.001** |
|  | imp18 | 4.59 | **0.0341** | 20.28 | **1.57E-05** | 3.7 | **0.0275** | 4.43 | **0.014** | 0.73 | 0.3935 |
|  | imp19 | 0.1 | 0.7477 | 34.79 | **3.52E-08** | 4.67 | **0.0111** | 2.69 | 0.072 | 0.27 | 0.604 |
|  | imp24 | 0.01 | 0.9266 | 18.73 | **3.15E-05** | 1.49 | 0.2297 | 7.08 | **0.0012** | 5.08 | **0.026** |
|  | imp23 | 12.45 | **0.0006** | 0.67 | 0.4158 | 2.64 | 0.0756 | 6.48 | **0.0021** | 8.26 | **0.0048** |
|  | imp27 | 13.35 | **0.0004** | 0.25 | 0.6199 | 3.95 | **0.0218** | 2.96 | 0.0557 | 0.94 | 0.3347 |
|  | imp28 | 5.64 | **0.0191** | 1.3 | 0.2558 | 1.92 | 0.1508 | 7.48 | **0.0009** | 0.66 | 0.4197 |

Traits are grouped according to their maximal weights in the respective factors (Table 3). F, Fisher’s test; p, the significance of effects of independent variables, including X, the X chromosome; Aut, the autosomes; ♂P, the paternal genotype; and ChrY*Aut, a combined effect of the Y chromosome and autosomes. HC, a group of the traits that were not incorporated in the factor structures with weights higher than |0.5|. Significance values р < 0.05 are in bold. In each group of traits determining the respective factor structure, the lowermost row shows the estimated effects of the independent variables on the given factor.

Suppl. Tabl.5. Permutational ANOVA to separate the genotypes into groups homogeneous in trait values

| *F##* | *Signs* |  | *X_Y (epist)* | *AUT (add)* | *X_P + AUT (dom)_P* | *Y→AUT (dom. epist)+Y* | *AUT (add)*_*P+Y_P* | *Y→AUT (rec.epist)+ X→AUT (rec.epist)* | *X→AUT (dom. epist) +X+AUT (dom)* |
| --- | --- | --- | --- | --- | --- | --- | --- | --- | --- |
| F1 | imp29 | Prob. | 0.355 | 0.160 | 0.508 | 0.966 | 0.217 | 0.339 | 0.115 |
|  |  | Dom. | n.s. | n.s. | n.s. | n.s. | n.s. | n.s. | n.s. |
|  | imp31 | Prob. | 0.351 | 0.804 | 0.012 | 0.150 | 0.091 | 0.844 | 0.051 |
|  |  | Dom. | n.s. | n.s. | +* sD^Vi^_Aut_ | n.s. | n.s. | n.s. | n.s. |
|  | imp33 | Prob. | 0.185 | 0.039 | 0.000 | 0.050 | 0.002 | 0.582 | 0.002 |
|  |  | Dom. | n.s. | n.s. | + D^Vi^_Aut_ | n.s. | +* D^Vi^_Aut, (X?)_ | n.s. | +* sD^Vi^_Aut_ |
| F1* |  | Prob.* | 0.493 | 0.476 | 0.473 | 0.136 | 0.302 | 0.638 | 0.304 |
|  |  | Dom.* | n.s. | n.s. | n.s. | n.s. | n.s. | n.s. | n.s. |
| F2 | imp30 | Prob. | 0.007 | 0.160 | 0.000 | 0.480 | 0.000 | 0.011 | 0.004 |
|  |  | Dom. | + | n.s. | + D^Lu^_X-chr_ | n.s. | + P_Y_ | + ID | +* sD^Lu^_X-chr*_ |
|  | imp32 | Prob. | 0.157 | 0.078 | 0.000 | 0.023 | 0.000 | 0.062 | 0.000 |
|  |  | Dom. | n.s. | n.s. | + sD^Lu^_X-chr_ | n.s. | *_(X-chr)_ | n.s. | +* |
|  | imp34 | Prob. | 0.103 | 0.000 | 0.000 | 0.025 | 0.000 | 0.025 | 0.000 |
|  |  | Dom. | n.s. | D^Vi^ | +* | n.s. | + D^Vi^_Aut_ | +* sD^Lu^_Y-chr_ | + sD^Vi^_Aut_ |
|  | beta* | Prob. | 0.001 | 0.007 | 0.000 | 0.331 | 0.000 | 0.000 | 0.003 |
|  |  | Dom. | + | ID | + D^Lu^_X-chr_ | n.s. | + D^Lu^_Aut_ | + D^Lu^_Y-chr_ | + ID |
| F2* |  | Prob.* | 0.113 | 0.207 | **0.000** | **0.038** | **0.000** | 0.057 | **0.000** |
|  |  | Dom.* | n.s. | n.s. | + sD^Lu^_X-chr_ | + | *_(X-chr)_ | n.s. | +*(sD^Vi^_X-chr*_) |
| F3 | imp4 | Prob. | 0.008 | 0.000 | 0.077 | 0.675 | 0.022 | 0.000 | 0.051 |
|  |  | Dom. | + | D^Lu^ | n.s. | n.s. | + D^Lu^_Aut_ | + D^Vi^_X-chr_ | n.s. |
|  | imp6 | Prob. | 0.000 | 0.000 | 0.015 | 0.446 | 0.000 | 0.000 | 0.000 |
|  |  | Dom. | + | ID | +* sD^Vi^_Aut_ | n.s. | + P_Y_ | + D^Vi^_X-chr_ | + D^Lu^_X-chr*_ |
|  | imp14 | Prob. | 0.000 | 0.000 | 0.074 | 0.020 | 0.000 | 0.000 | 0.000 |
|  |  | Dom. | + | ID | n.s. | n.s. | + P_Aut_ | + D^Vi^_X-chr_ | + D^Vi^_Aut_ |
|  | imp16 | Prob. | 0.000 | 0.000 | 0.006 | 0.320 | 0.000 | 0.000 | 0.000 |
|  |  | Dom. | + | ID | +* | n.s. | + P_Aut_ | + D^Vi^_X-chr_ | + ID |
|  | imp21 | Prob. | 0.246 | 0.000 | 0.279 | 0.722 | 0.058 | 0.000 | 0.049 |
|  |  | Dom. | n.s. | D^Lu^ | n.s. | n.s. | n.s. | +* sD^Vi^_X-chr_ | n.s. |
|  | imp25 | Prob. | 0.000 | 0.000 | 0.000 | 0.311 | 0.000 | 0.000 | 0.000 |
|  |  | Dom. | + | ID | + D^Lu^_X-chr_ | n.s. | + P_Aut_ | + D^Vi^_X-chr_ | + ID |
| F3* |  | Prob.* | **0.001** | **0.000** | **0.000** | 0.285 | **0.000** | **0.000** | **0.000** |
|  |  | Dom.* | + | D^Lu^ | +* | n.s. | + P_Y_, D^Lu^_Aut_ | +* sD^Vi^_X-chr_ | +* sD^Lu^_X-chr*_ |
| F4 | imp11 | Prob. | 0.100 | 0.021 | 0.000 | 0.000 | 0.000 | 0.019 | 0.000 |
|  |  | Dom. | n.s. | D^Lu^ | +* | + | + P_Y_ | + ID | + D^Lu^_X-chr*_ |
|  | imp20 | Prob. | 0.041 | 0.000 | 0.000 | 0.001 | 0.000 | 0.001 | 0.000 |
|  |  | Dom. | + | ID | +* | + | + P_Y_ | + D^Vi^_X-chr_ | + D^Lu^_X-chr*_ |
| F4* |  | Prob.* | 0.697 | 0.489 | **0.000** | **0.000** | **0.000** | 0.738 | **0.000** |
|  |  | Dom.* | n.s. | n.s. | +* | + | + P_Y_ | n.s. | +* sD^Lu^_X-chr*_ |
| F5 | imp8 | Prob. | 0.431 | 0.000 | 0.347 | 0.739 | 0.000 | 0.696 | 0.000 |
|  |  | Dom. | n.s. | D^Vi^ | n.s. | n.s. | + D^Vi^_Aut_ | n.s. | + D^Vi^_Aut_ |
|  | imp10 | Prob. | 0.000 | 0.000 | 0.000 | 1.000 | 0.000 | 0.000 | 0.000 |
|  |  | Dom. | + | D^Lu^ | +* | n.s. | +* D^Lu^_Aut_ | + D^Vi^_X-chr_ | + D^Lu^_X-chr_ |
|  | imp13 | Prob. | 0.097 | 0.278 | 0.000 | 0.260 | 0.023 | 0.964 | 0.011 |
|  |  | Dom. | n.s. | n.s. | + D^Vi^_Aut_ | n.s. | +* D^Lu,Vi^_Aut_ | n.s. | +* |
|  | imp15 | Prob. | 0.002 | 0.031 | 0.368 | 0.177 | 0.013 | 0.020 | 0.031 |
|  |  | Dom. | + | sD^Vi^ | n.s. | n.s. | +* D^Vi^_Aut_ | +* sD^Lu^_Y-chr_ | +* sD^Vi^_Aut_ |
| F5* |  | Prob.* | **0.000** | **0.000** | 0.043 | 0.025 | **0.001** | **0.000** | 0.051 |
|  |  | Dom.* | + | +* | n.s. | + | +* D^Lu^_Aut_ | + ID | n.s. |
| F3, F6 | imp2 | Prob. | 0.001 | 0.000 | 0.000 | 0.121 | 0.000 | 0.001 | 0.000 |
|  |  | Dom. | + | sD^Lu^ | + ID | n.s. | *_X-chr_ | + D^Vi^_X-chr_ | +* |
| F6 | alpha | Prob. | 0.717 | 0.000 | 0.000 | 0.016 | 0.000 | 0.189 | 0.000 |
|  |  | Dom. | n.s. | D^Vi^ | + D^Vi^_Aut_ | + | + D^Vi^_Aut_ | n.s. | +* sD^Vi^_Aut_ |
| F6* |  | Prob.* | 0.478 | **0.000** | **0.000** | **0.000** | **0.000** | 0.857 | **0.000** |
|  |  | Dom.* | n.s. | D^Vi^ | + D^Vi^_Aut_ | + | +D^Vi^_Aut_ (*_X-chr_) | n.s. | + ID |
| F7 | imp5 | Prob. | 0.026 | 0.011 | 0.092 | 0.078 | 0.043 | 0.044 | 0.382 |
|  |  | Dom. | + | sD^Lu^ | n.s. | n.s. | n.s. | n.s. | n.s. |
|  | imp7 | Prob. | 0.804 | 0.002 | 0.122 | 0.150 | 0.017 | 0.036 | 0.003 |
|  |  | Dom. | n.s. | sD^Vi^ | n.s. | n.s. | + D^Vi^_Aut_ | n.s. (D^Vi^_X-chr_) | + D^Vi^_Aut_ |
|  | imp9 | Prob. | 0.001 | 0.001 | 0.220 | 0.023 | 0.001 | 0.005 | 0.004 |
|  |  | Dom. | + | ID | n.s. | n.s. | + D^Vi^_Aut_ | + ID | + ID |
|  | imp17 | Prob. | 0.440 | 0.000 | 0.010 | 0.001 | 0.000 | 0.547 | 0.000 |
|  |  | Dom. | n.s. | D^Vi^ | +* sD^Lu^_X-chr_ | + | + D^Vi^_Aut_ | n.s. | + D^Vi^_Aut_ |
| F7* |  | Prob.* | 0.542 | **0.000** | **0.045** | **0.031** | **0.002** | **0.028** | **0.001** |
|  |  | Dom.* | n.s. | sD^Vi^ | n.s.(sD^Lu^_X-chr_) | + | +* D^Vi^_Aut_ | +* | + D^Vi^_Aut_ |
| CH | imp3 | Prob. | 0.000 | 0.000 | 0.811 | 0.009 | 0.000 | 0.000 | 0.123 |
|  |  | Dom. | + | D^Lu^ | n.s. | + | + D^Lu^_Aut_ | + D^Vi^_X-chr_ | n.s. |
|  | imp18 | Prob. | 0.000 | 0.000 | 0.222 | 0.008 | 0.009 | 0.000 | 0.085 |
|  |  | Dom. | + | D^Lu^ | n.s. | + | + P_Aut_ | + D^Vi^_X-chr_ | n.s. |
|  | imp19 | Prob. | 0.000 | 0.000 | 0.302 | 0.000 | 0.002 | 0.000 | 0.009 |
|  |  | Dom. | + | ID | n.s. | + | + D^Vi^_Aut_ | + ID | + D^Vi^_Aut_ |
|  | imp23 | Prob. | 0.385 | 0.004 | 0.033 | 0.412 | 0.002 | 0.000 | 0.002 |
|  |  | Dom. | n.s. | D^Vi^ | n.s. | n.s. | +* D^Vi^_Aut_ | +* | + ID |
|  | imp24 | Prob. | 0.010 | 0.004 | 0.000 | 0.000 | 0.004 | 0.003 | 0.016 |
|  |  | Dom. | + | D^Vi^ | + D^Vi^_Aut_ | + | + D^Vi^_Aut_ | + ID | +* sD^Vi^_Aut_ |
|  | imp26 | Prob. | 0.055 | 0.000 | 0.239 | 1.000 | 0.145 | 0.000 | 0.120 |
|  |  | Dom. | n.s. | D^Lu^ | n.s. | n.s. | n.s. | + D^Vi^_X-chr_ | n.s. |
|  | imp27 | Prob. | 0.306 | 0.000 | 0.054 | 0.253 | 0.004 | 0.000 | 0.002 |
|  |  | Dom. | n.s. | D^Lu^ | n.s. | n.s. | +*_X-chr_ | +* sD^Vi^_X-chr_ | + D^Lu^_X-chr*_ |
|  | imp28 | Prob. | 0.020 | 0.538 | 0.000 | 1.000 | 0.002 | 0.042 | 0.067 |
|  |  | Dom. | + | n.s. | + D^Lu^_X-chr_ | n.s. | + D^Lu^_Aut_ | n.s. | n.s. |
| FDR | 0.05 | | 0.023 | 0.037 | 0.026 | 0.010 | 0.040 | 0.031 | 0.034 |
|  | 0.01 | | 0.003 | 0.006 | 0.004 | 0.001 | 0.007 | 0.005 | 0.006 |

(+), a significant effect of a hereditary factor that is in linear relationship with the given genotype classes; (+*), a significant effect of a hereditary factor that is in nonlinear relationship with the given genotype classes; D^Vi (Lu)^, dominance of the *D. virilis* (*D. lummei*) phenotype in genotypes with the intermediate value of the indicator variable; P_Y(Aut)_, an epigenetic effect of the male parent identity on the Y chromosome (autosomes); the subscripts Aut, X-chr, and Y-chr indicate that the autosome set, X chromosome, or Y chromosome determines phenotype dominance; F#, latent trait number as in Table 3; F#*, permutation test values for the factors (latent traits) used as variables; CH, highly characteristic trait; FDR, false discovery rate for the primary traits.
